# Nicotine Influence on Vascular and Neurocognitive Function with In-utero Electronic Cigarette Exposure

**DOI:** 10.1101/2025.02.13.638202

**Authors:** Amber Mills, Deborah Corbin, Duaa Dakhallah, Paul D. Chantler, I. Mark Olfert

## Abstract

Emerging studies find arteriolar dysfunction in offspring with in-utero electronic cigarette (Ecig) exposure, but the long-term effect on offspring’s cerebrovascular vascular and neurocognitive health is poorly understood. Ecigs provides a unique opportunity to directly evaluate the contributions of inhaled nicotine from the vehicle e-liquid – which was not possible with traditional cigarettes. Moreover, many Ecigs have variable power settings, which can alter the toxicity of the aerosol cloud produced. We hypothesize maternal vaping at different wattages will have variable effects on cerebrovascular function in the offspring, and that these effects would be independent of nicotine. We used time-mated female Sprague-Dawley rats with Ecig exposure from gestation day (GD)2-21. We studied male and female offspring for vascular and neurocognitive function at 1-, 3-, 6- and 12-months of age. We found that, both sexes, offspring with in-utero exposure (at 5w and 30w Ecig conditions) exhibited impaired middle cerebral artery (MCA) reactivity. While the magnitude of impairment was greater at higher that lower watts, Ecig at 5-watts still exhibited significant impairments in MCA function (suggesting the harm threshold for blood vessels is very low). Vascular dysfunction was evident with or without nicotine in the e-liquid, but nicotine exposure resulted in short-term memory deficits, evidence of neuronal damage, and increased astrocyte interaction with endothelial cells in 6- and 12-month-old offspring. We also observed altered expression of clock genes and antioxidant signaling pathways, along with a decrease in sirtuin-1 expression, decreased ratio of beta-amyloid Aꞵ 42/40 protein expression, and increased in NOX1, which are consistent with redox imbalance, neuroinflammation, and advancing cellular senescence. These preclinical data provide evidence suggesting that in utero exposure to Ecigs from maternal vaping can be expected to adversely affect the brain health of offspring in their adult life and that neurocognitive outcomes are worsened with exposure to nicotine.

**Graphical Abstract:** 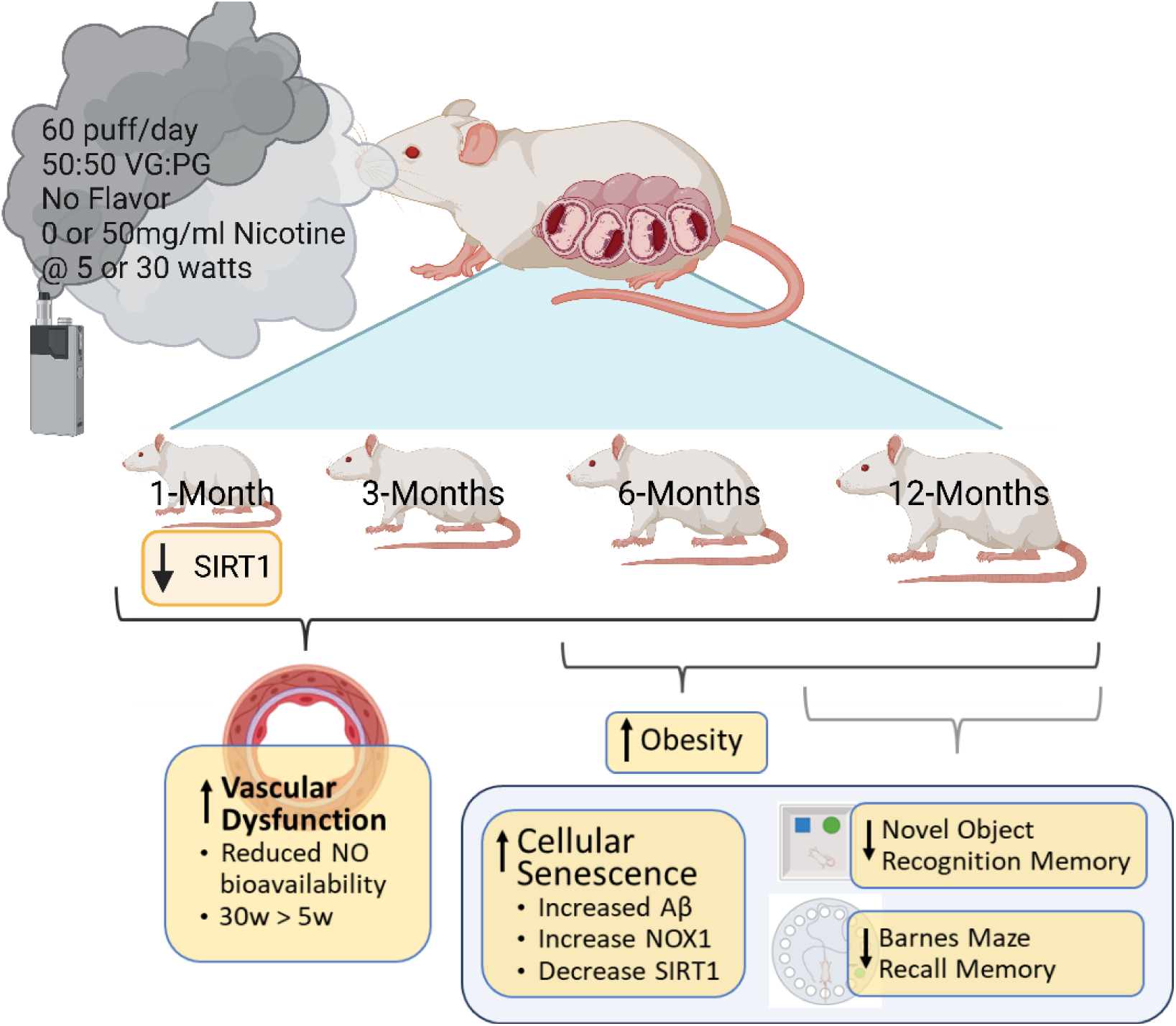

Created in BioRender

## INTRODUCTION

Electronic cigarettes (Ecigs) have grown in popularity based on the assumption and advertising that they are ‘safer’ than traditional cigarettes, but the fact remains that Ecigs still contain nicotine and a wide array of harmful toxins. For example, Ecig aerosol contains carbonyl compounds, reactive aldehydes, and metals [1–3], some of which, like formaldehyde, are known to be carcinogenic [4–7]. These carcinogens can alter basic cell processes (e.g., autophagy, etc.) and lead to cellular damage [8–10]. Direct E-cig exposure in mice has been shown to disrupt clock genes, leading to DNA damage and early cellular senescence [11, 12]. There is recent evidence showing maternal use during pregnancy results in cerebrovascular and cardiovascular impairment to the maternal-fetal dyad [13–23]. Importantly, as Ecigs have grown in popularity, the concentration of nicotine use (compared to cigarettes) has also increased [24]. This increase is concerning since nicotine use during pregnancy has been linked in both animals and humans to poor fetal outcomes (such as low birth weight, small for gestational age, neurocognitive deficits, and more)[25–29]. Yet, it is interesting to note that even Ecig use without nicotine is reported to result in vascular dysfunction [22, 23], and the dysfunction from Ecigs is comparable to vascular deficits from smoking [30–33]. Thus, the effects of inhaling nicotine during pregnancy on offspring health remain complex and require further investigation.

The direct effects of nicotine include blood vessel constriction, which can decrease tissue blood flow and alter nutrient supply and cellular health [34]. Nicotine can also cause a shift in the circadian rhythm via nicotinic acetylcholine receptors (nAChRs) in the suprachiasmatic nucleus (SCN, an area known as the master circadian clock)[35]. Disruption in circadian clock rhythm is reported to increase cellular aging, impact behavior, hormone balance, and can be an early warning sign for neurodegenerative diseases such as Alzheimer’s [36–39]. The base e-liquid in Ecigs (i.e., vegetable glycerin, VG; and propylene glycol, PG) has also been shown to trigger changes in clock gene expression [40]. Moreover, exposure to environmental tobacco smoke can interfere with the molecular clock by disrupting key genes that regulate the circadian clock (such as reduced expression of BMAL1 and downstream SIRT1 signaling) in both mice and patients with COPD [41, 42]. SIRT1 is of particular interest because it is a nicotine adenine dinucleotide (+) [NAD+]-dependent histone deacetylase that regulates many cellular processes, such as metabolism, inflammation, aging and more. For example, SIRT1 regulates glucose and lipid metabolism by deacetylating substrates. SIRT1 is downregulated by autophagy during senescence and aging, and SIRT1 is essential for cognitive function and synaptic plasticity [43]. While there is strong evidence that Ecigs create the same vascular dysfunction as smoking [14], and smoking is widely recognized as a major risk factor for accelerated cellular and function decline in humans, including vascular function [44], little is still known about the long-term cellular effects related to key molecular regulators, like SIRT1, and clock genes (regulating circadian rhythm) in offspring with in utero exposure to Ecigs from maternal vaping (with or without nicotine). Thus, we sought to examine the effects of maternal Ecig use during pregnancy (with or without nicotine) on cerebral vascular and neurocognitive health of offspring in their postnatal adult life.

## MATERIALS AND METHODS

### Ethical Approval

All procedures were conducted with approval from West Virginia University Animal Care and Use Committee (IACUC) approval (#16-05003053), and to establish principles and regulations for animal experimentation [45]. Surgical procedures were terminal and performed under deep anesthesia (inhaled isoflurane) to minimize pain or discomfort.

### Study design and exposure system

Female (200-250 g) and male (250-300 g) Sprague-Dawley rats were purchased for breeding (Charles River, Wilmington, MA) and housed in an AAALAC-certified vivarium facility. Rodents were provided standard rat chow and reverse osmosis drinking water and kept on a 12- h:12-h day/night cycle, throughout the study at West Virginia University. Animals were allowed to acclimate to the new facility for at least 7 days before breeding. Before mating, vaginal smears were obtained at least for three consecutive days to determine the female’s estrous cycle, and males were added during the start of estrous. Males were housed with females around 5 pm and removed the following morning between 8-10 am to correspond with their active cycle. After which a vaginal smear was obtained to verify the presence of sperm or a vaginal plug as evidence of pregnancy and establishment of gestational day (GD)0. Dams were then randomly assigned to receive the following exposures:

1: Ecig aerosol with no nicotine (Ecig0) at 5-Watts (Ecig0-5W, n=5) 2: Ecig aerosol with no nicotine at 30-Watts (Ecig0-30W, n=5)
3: Ecig aerosol with nicotine at 5-Watts (Ecig50-5W, n=5)
4: Ecig aerosol with nicotine at 30-Watts (Ecig50-30W, n=5)
5: Ambient air (control, n = 6)

Air controls were handled at the same time as the exposed animals and were placed in the same exposure room to replicate the stress of handling and transport but were only exposed to ambient air.

Maternal exposure began on either gestational day (GD)2 until GD21. The exposure consisted of 60 puffs/day (1 puff every 3 min over 90 min), 5 days/week for 3 weeks (GD2-GD21) using a whole-body chamber. Puff duration was set at 5 s with an inhalation draw rate of ∼1 LPM and a 20L/min bias flow of air was used to continuously flush the chamber throughout the exposure which provided an episodic exposure pattern similar to the episodic vaping behavior in humans [46]. E-liquid consisted of a mixture of 50:50 vegetable glycerin (VG): propylene glycol (PG) with no flavoring. VG was purchased from Sigma (#2143-01, Avantor, J.T. Baker) and PG from Fisher Scientific (#P355-1). A 1.2 Ω Joyetech atomizer was used for 5W, whereas a 0.5 Ω Joyetech atomizer was employed for 30W exposure.

### Aerosol Analysis

Chamber aerosol concentrations were obtained and recorded in real-time throughout each exposure conditions using a Casella MicroDust Pro Monitor (Model CEL-712). Offline assessments for aerosol size distribution were also made using electrical low-pressure impactor (ELPI+, Dekati Ltd) under the wattage conditions used in this study (i.e. 5 and 30 watts) and have been previously reported [15].

### Measurements of Vascular Reactivity

Vascular function in offspring was studied at 1- 3- and 6- and 12-months of age, where 1 male and 1 female from each litter (at each time point) were euthanized to assess the middle cerebral artery (MCA) reactivity using pressure myography (Scintica Inc. London, ON, Canada). Rats were anesthetized by 5% isoflurane inhalation for induction and maintained on a plane of anesthesia at 2-3% until euthanized. Euthanasia occurred by exsanguination via cardiac puncture. After blood collection, the vascular system was flushed with phosphate buffer saline (PBS) solution, and the brain quickly and carefully was removed to obtain the MCA vessel. The MCA was isolated from Circle of Willis, cleaned and placed into a microvessel chamber filled with cold physiological salt solution (PSS; 4°C). Both ends of the isolated MCA were cannulated within a chamber that allowed the lumen and exterior of the vessel to be perfused and superfused, respectively, with heated PSS (37°C) from separate reservoirs. PSS was equilibrated with a 21% O2, 5% CO2, and 74% N2 gas mixture and had the following composition (mM): 119 NaCl, 4.7 KCl, 1.17 MgSO4, 1.6 CaCl2, 1.18 NaH2PO4, 24 NaHCO3, 0.026 EDTA, and 5.5 glucose.

Following cannulation, MCAs were extended to their in-situ length and equilibrated to ∼70 mmHg [32]. Then the vessel was challenged using 10^-4^ M serotonin to check viability; thereafter, vessel reactivity was assessed in response to increasing concentrations of an endothelial-dependent dilator (EDD) (acetylcholine, ACh; 10^-9^ M – 10^-4^ M), an endothelial-independent dilator (EID) (sodium nitroprusside, SNP; 10^-9^ M – 10^-4^ M), and a vasoconstrictor agent (serotonin, 5-HT; 10^-9^ M – 10^-4^ M). Additionally, MCA responses to ACh were measured following acute incubation (30 minutes) with nitro-L-arginine methylester (L-NAME, 10^-4^ M; an inhibitor of NO synthase, Sigma Aldrich) to assess contributions of nitric oxide (NO); TEMPOL, a superoxide dismutase to better understand the implication of oxidative stress; and Febuxostat, a xanthine oxidase inhibitor to evaluate the effects of superoxide and hydrogen peroxide on MCA reactivity [33]. A video dimension analyzer connected to an inverted microscope was used to measure lumen diameter. The MCA has endogenous smooth muscle tone therefore pre-constriction prior to assessing dilator responses is not required. Thus, dilation and constriction are assessed by subtracting the starting lumen diameter before each treatment and chambers were washed between each drug dose response.

After completion of vascular reactivity measurements, we also examined the active and passive myogenic responses in MCA’s of 12-month-old male and female offspring. Vessels were exposed to each pressure point in a non-consecutive way for 5 min before each reading was recorded. For active myogenic response, inner and outer diameter curves were obtained in the PSS with Ca^2+^ to observe the vessels’ contractile properties and then in Ca^2+^-free PSS to evaluate the vessels’ passive properties. Additional calculations of arteriolar wall mechanics are based on previously reported methodology [23, 47].

### Neurocognitive and behavioral testing

We used several tests to assess rodent cognitive function and behavior. Y-maze testing was performed at ∼8 months of age, while novel object recognition (NOR) and Barnes maze between 11-12 months of age. Open field testing was conducted as repeated measures (using the same animals) at 3-, 6- and 12-months of age. Methods employed for each test is as follows:

#### Novel object recognition

Two pairs of identical objects and the black Plexiglas arena were previously validated to ensure the animals had no preference for this experiment. Validation of the objects occurred with a subset of rats (n=20) similar in age to the experimental rats and contained males and females but were not used for the study. For validation, both sets of objects were introduced and then the time spent exploring each object in a 5-minute trial was recorded. The exploration time of Object A / Object B was recorded as the bias ratio was set to anything less than or greater than 0.8-1.2. Only the 2 sets of objects that were closest to 1.0 (i.e. equally attractive) were used.

For experimental animals, one pair of identical objects were placed in the arena before adding the 12-month-old rats to habituate for 15 min on day 1. At the end of each trial rats were placed back in their home cages and left rest for 1 h. In between each animal all objects and the area were cleaned with 70% ethanol to remove any olfactory cues. Animals were again placed in the area with the similar objects for an additional 5-mintues.

The following day, one of the familiar objects was replaced by a novel object, and animals were allowed to explore the arena for 5 min. Either the right or left object was replaced randomly for each animal with the novel object. Object interaction was automatically recorded with a video tracking system (AnyMaze Systems). Time spent interacting was recorded in sec for the 5-minute novel trial. Using the AnyMaze program, representative track plots of the final trial were created, and time interacting with the familiar and novel object was recorded.

#### Barnes Maze

The maze consisted of a tan platform (130 cm in diameter, 91 cm height) with 20 holes (10 cm in diameter) located 2.5 cm from the perimeter (San Diego Instruments) grey escape box (EB) was placed under one of the holes, and 19 false grey target boxes were placed under all other holes. 12-month-old rats were brought into the testing room 1 h before the experiment for acclimation. All rats were trained for five trials from day 0 to 5 then a probe trial was performed. The probe trial consisted of the removal of the EB to be replaced with another false target box. Therefore, there is no longer an escape hole for the rats to enter. Then the rats had a three-day reversal retrial where the EB was placed on the opposite side of the maze, and another probe trial was performed after the reversal. All rats began in the center of the maze with a partition box preventing them from seeing the maze for 30 s before the trial began. During training sessions, the rat was allowed to freely explore the maze until either entering the EB or after 2 min elapsed. Once the rat entered the EB, it was left there for 30 s before returning to their home cage. If the rat did not enter the EB by itself, it was gently guided to and allowed to stay in the EB for 30 s. After the training session, rats were tested with 5-trial over 5 consecutive days. Testing was similar to training, but if after 2 min the rat did not find the escape box, it was directly returned to its home cage. For the acquisition trial, path efficiency was recorded and tracked to see each animal’s progress over the 5-days. During the reversal, an average of all the animals in each group was used to make a heat map to show the time spent. Lastly, the number of animal interaction with the EB and 19 false boxes were recorded.

#### Y-maze test

The Y maze is a three-arm maze, made of black Plexiglas with equal angles between all arms. 8-month-old offspring were individually tested by placing them within the same arm of the maze (labeled A) and allowing them to move freely throughout the three different arms of the maze over an 8 min period. The sequence and entries in each arm were recorded, and alternation was determined from successive consecutive entries to the three different arms on overlapping triads in which all arms were represented. For example, a sequence of entries such as ACB where all three arms were entered would be counted as a successful alternation, and ABA would be an unsuccessful alternation because an arm was repeated. The percentage of alternations per total entry was then calculated [48] along with an average heat map to show time spent in the Y maze.

#### Open field (OF)

To evaluate anxiety-like behaviors at the same time points, all offspring in the litter were assessed with OF testing. The offspring were individually placed in a photobeam activity system open field chamber for 30-minutes after a 1-hour room acclimation. During that time the animals were not disturbed and allowed to move freely. Both X and Y plane lasers within the chambers were used to track movement. Changes in central tendency (i.e. the percentage of time spent in the center of the open field compared to the percentage of time spent in the periphery), locomotor activity, and fecal count are reported. For fecal counts, droppings that were not intact or were not solid were excluded. In every test, males were run first then females and chambers were thoroughly cleaned to remove any olfactory cues before the start of another test.

### Protein expression in cerebral tissue

The level of Sirtuin-1 (SIRT1, NOVUS Biologicals, Catalog #NBP2-80302) and NADPH oxidase (NOX1, NOVUS Biologicals, Catalog #NBP2-76748) was assessed in 1- and 12-month offspring according to manufacturer instructions. Cerebral cortex homogenates ∼200mg of tissue was used with ∼100mg used per well to obtain duplicate assessment. Levels of Amyloid Beta (Aꞵ)1-42, Aꞵ1-40 (NOVUS Biologicals, Catalog #NBP2-69916, Fisher Scientific, Catalog #50- 149-8811, respectively) were also evaluated in 12-month offspring using cerebral homogenates keeping the same tissue volumes as above.

### Gene expression of clock genes

RNA was isolated from cortex and cerebellum samples using (QIAGEN RNeasy Lipid Tissue Mini Kit, Catalog #74804) according to the protocol, and RNA quality was assessed using Nanodrop Spectrophotometer. cDNA synthesis was performed on 1ug of RNA using SuperScript™ III First-Strand Synthesis SuperMix for qRT-PCR (Thermofisher, Catalog # 11752050). SYBR Green master mix (Life Technologies, Grand Island, NY) and verified primers (IDT Inc. Coralville, Iowa) following manufacturer’s instructions. Reactions were carried out on the Quant Studio 6 Flex. mRNA expression was normalized to adenylate cyclase-associated protein 1 (CAP1) or ribosomal protein L4 (RPL4).

### Histology of Coronal Section

Free-floating coronal sections (30 um) were obtained via cryosectioning and underwent silver staining with FD NeuroSilver Kit II (CAT# PK301A) according to the manufacturer’s directions. Sections were scored by an expert blinded to the group identity. Immunohistochemistry was also performed on separate sections to evaluate astrocytes (via glial fibrillary acidic protein, GFP, Invitrogen catalog #A11008) and endothelial cells (using tomato lectin Vector Labs catalog #DL-1177). Here, tissue sections were washed with 0.1M phosphate-buffered saline (PBS), blocked with 10% normal goat serum, and incubated overnight in the primary antibody. The following day, tissue was incubated in secondary antibody (i.e. goat anti-rabbit secondary GFP) and Tomato lectin antibody for 2 h, washed, and mounted. Fluorescent sections were imaged using the MIF Zeiss 710 Confocal microscope at 20X for all images (astrocytes excitation at 488 nm and endothelial cells at 594 nm).

### Collection of Extracellular Vesicles

Plasma from offspring at 12-month-old of age was centrifuged at 1,500 rpm, 4 °C, for 10 min to separate plasma and cell debris. The supernatant was removed and placed in a new tube. Plasma EVs were purified by centrifugation at 16,500 rpm, 4 °C for 1 h. EVs pellet was washed with sterile and filtered PBS and resuspended in 200 µl of PBS. Thereafter, 1 µl was taken and diluted in 1 mL of filtered 1XPBS (without Ca and Mg) for analyses of EV size and quantity (Malvern Panalytical Nanosight NS300). These samples were later examined using the Exo-check antibody array (No. EXORAY200A-4, Thermo Fisher Scientific) using 50 µg of protein to confirm the presence/identity that particles were EVs.

### Data and Statistical Analyses

All data are presented as mean ± SE, except when noted. Vessel reactivity to changes in drug dose concentration was first analyzed by repeated-measure two-way analysis of variance (rANOVA). If main effects were observed, a post-hoc (Tukey) test was used to determine differences between conditions (i.e. time and exposure group). Maximal responses were independently analyzed using one-way analysis of variance (ANOVA) to evaluate exposure group differences. Tukey post-hoc test was used to determine differences between groups, when appropriate. In all cases, p ≤ 0.05 was taken to reflect statistical significance.

## Results

Ecig device settings and chamber conditions are reported in **Table 1**. We saw no difference in chamber temperature and humidity. However, as expected mass concentration of Ecig aerosol for 30W exposures was significantly greater compared to 5W exposures (p<0.05, **Table 1**).

**Table 1.** Chamber and Ecig Device Conditions.

|  | **Air** | **Ecig0**<br>**5W †** | **Ecig50**<br>**5W** | **Ecig0**<br>**30W †** | **Ecig50**<br>**30W** |
| --- | --- | --- | --- | --- | --- |
| **<u>E-cig Device</u>** |  |  |  |  |  |
| Watts | - | $5 \pm 0.1$ | $5 \pm 0.2$ | $29 \pm 1$ | $30 \pm 0.7$ |
| Volts | - | $2.5 \pm 0.3$ | $2.6 \pm 0.1$ | $3.8 \pm 0.2$ | $3.9 \pm 0.1$ |
| Amps | - | $2.0 \pm 0.2$ | $2.1 \pm 0.1$ | $7.4 \pm 0.4$ | $7.5 \pm 0.2$ |
| **<u>Chamber</u>** |  |  |  |  |  |
| Temperature (C) | $68 \pm 2$ | $69 \pm 2$ | $69 \pm 1$ | $69 \pm 2$ | $69 \pm 1$ |
| Humidity (%) | $43 \pm 8$ | $31 \pm 12$ | $25 \pm 4$ | $28 \pm 8$ | $26 \pm 6$ |
| Aerosol Concentration (mg/m<sup>3</sup>) | - | $163 \pm 126$ | $231 \pm 125$ | **$917 \pm 382^*$** | **$1346 \pm 369^*$** |

† Data have been previously reported [82].

Maternal and pregnancy outcomes are reported in **Table 2**. There was no difference in the age of the dams at the time of pregnancy, the number of implantation sites, or the ratio of male to female offspring born. However, average litter size was significantly decreased with 30W exposure groups (i.e. Ecig0-30W and Ecig50-30W) compared to Air controls (p<0.05, **Table 2**). The number of reabsorption sites (# of implantation sites - # offspring born) was significantly increased in all Ecig groups (regardless of wattage or nicotine) litters compared to Air controls (p<0.05, **Table 2**).

**Table 2.** Litter Characteristics.

| **Litter Characteristics** |  |  |  |  |  |
| --- | --- | --- | --- | --- | --- |
|  | Air | Ecig0-5W† | Ecig50-5W | Ecig0-30W† | Ecig50-30W |
| Dams n= | 11 | 6 | 7 | 8 | 6 |
| Age of dams (months) | 4.6 ± 1.6 | 4.0 ± 0.6 | 3.3 ± 1.0 | 3.8 ± 0.6 | 3.4 ± 0.5 |
| Average Litter Size (#) | 14 ± 1.6 | 12 ± 1.9 | 11 ± 2.7 | **8 ± 3.8\*** | **8 ± 4.2\*** |
| Implantation Sites (#) | 14 ± 2 | 14 ± 1 | 13 ± 1 | 13 ± 2 | 13 ± 2 |
| Reabsorption Sites (#) | 0.2 ± 0.3 | **5.3 ± 1.1\*** | **4.2 ± 3.7\*** | **6.3 ± 1.8\*** | **5.4 ± 2.6\*** |
| Male vs Female Ratio | 1.4 ± 0.5 | 0.8 ± 0.3 | 0.8 ± 0.6 | 1.0 ± 0.5 | 1.1 ± 0.7 |

† Data have been previously reported [82].

Offspring weight was recorded at 1-, 3-, 6-, and 12-months of age (**Figure 1**). Male (no nicotine) exposed offspring gained significantly more weight compared to controls at 6- and 12- months (p<0.05). Females only surpassed Air control weights in the Ecig0-30W group at 12- months of age (p<0.05, **Figure 1A**). Both male and female offspring (with in-utero nicotine exposure) gained significantly more weight compared to Air controls at 6- and 12-months of age (p<0.05, **Figure 1B**), except for female Ecig50-5W (p=0.07).

**Figure 1:**
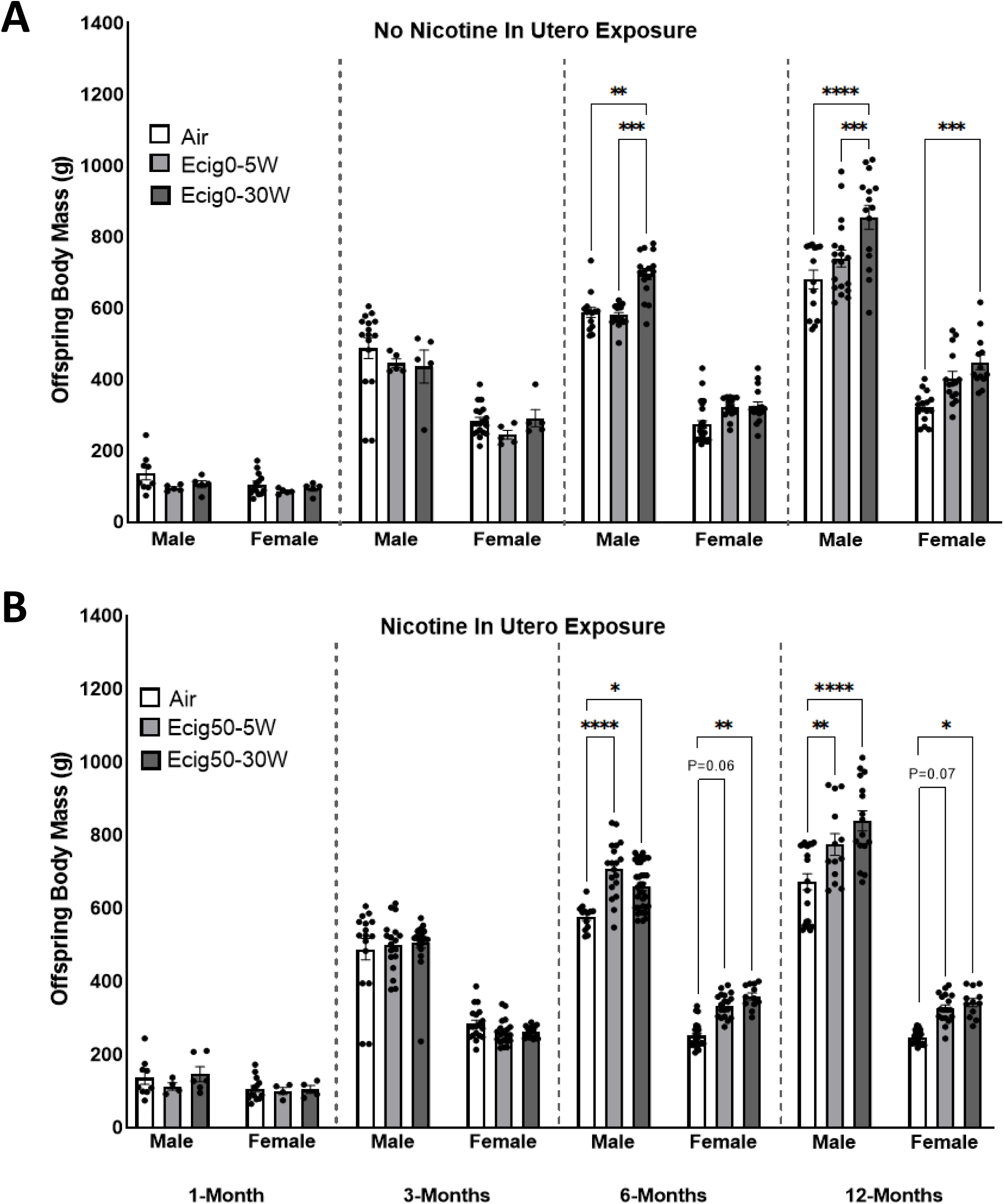
Offspring Weight at 1, 3, 6, and 12-Months of age separated by sex and nicotine (**A**-No Nicotine, **B**-Nicotine) exposure Mean ± SE.

MCA reactivity of offspring was assessed with ACh, SNP and 5-HT and reported in **Figure 2**. For all Ecig groups regardless of offspring age, MCA response to ACh was significantly blunted compared to Air controls (p<0.05, **Figure 2A**). The severity of impairment was wattage dependent, with 30W-exposed offspring having greater blunting of MCA ACh responses compared to 5W groups (p<0.05). Lastly, the addition of nicotine did not change MCA response to ACh in either wattage group (**Figure 2A**). MCA response to SNP was less severe compared to ACh, and was mostly observed at younger ages (i.e. 1- and 3-months), with the exception for Ecig0-30W exposure seen at all ages (p<0.05, **Figure 2B**). The MCA contractile response to serotonin (i.e. 5-HT) was impaired compared to air in the Ecig offspring at 1- and 6-months old, whereas at 12-months of age, the offspring with nicotine exposure (Ecig50-5W and Ecig50-30W) had hyperconstrictor response (**Figure 2C**). Myogenic MCA responses were also measured and found to not be different (Supplement Figure 1).

**Figure 2:**
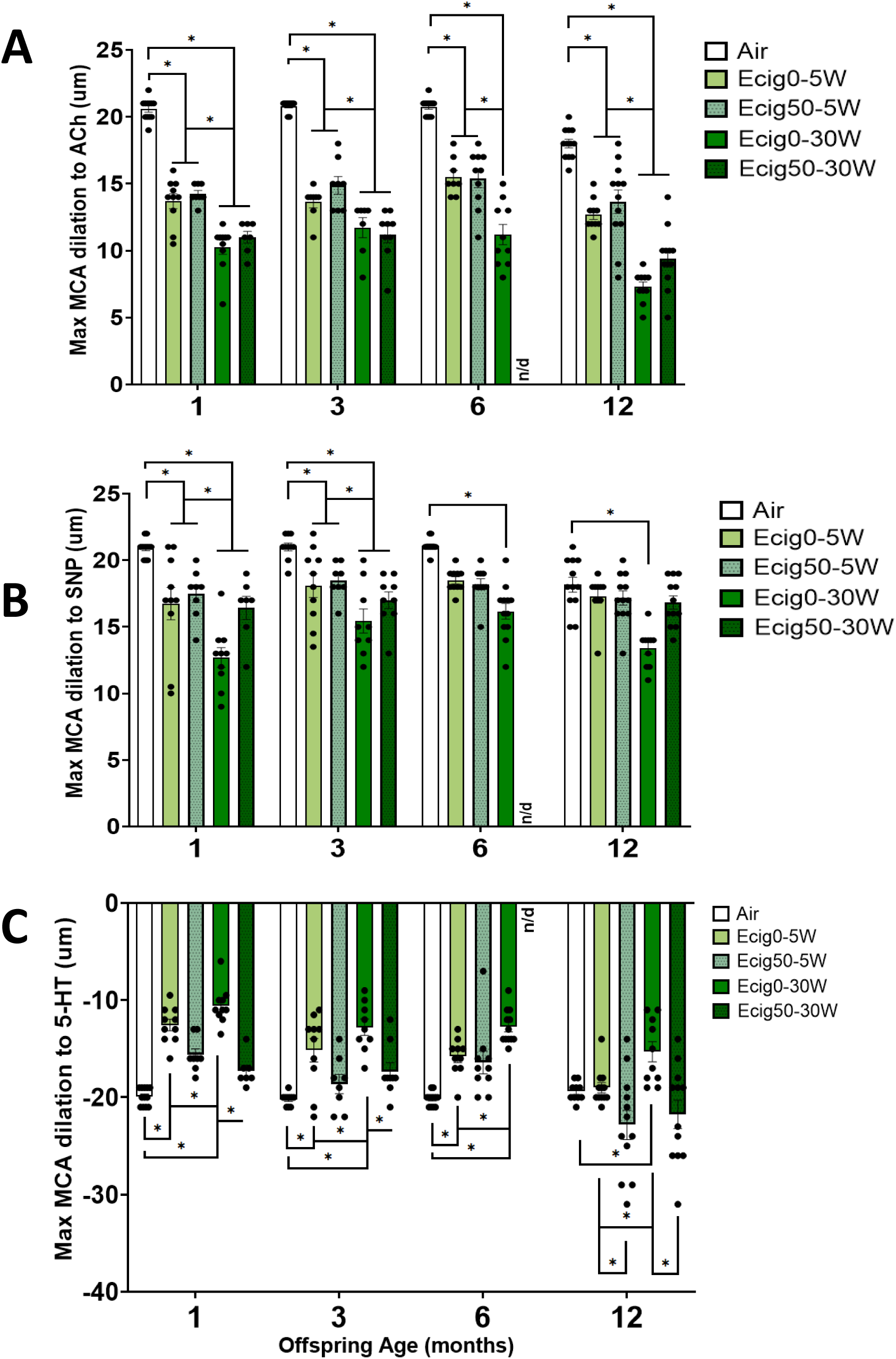
Middle Cerebral Artery (MCA) of 1-, 3-, 6-, and 12-months offspring maximal response to ACh to assess endothelial dependent dilation (EDD) **A**, SNP for endothelial-independent dilation (EID) **B**, (n = 5-6 dams/group sexes evenly distributed). Mean ± SE.

MCA responses to ACh after co-incubation with L-NAME show that inhibition of eNOS pathways significantly blunts, but does not fully eliminate, MCA reactivity. The impairment was not different based on the wattage used in non-nicotine exposure offspring, but did appear to be influenced with Ecig exposure including nicotine (**Figure 3A** no nicotine, **B** nicotine). The addition of TEMPOL improved MCA response to ACh in no nicotine offspring and also had significant improvement in all Ecig groups compared to ACh only, except for Ecig0-30W at 1-month of age (p<0.05, **Figure 4A**). All nicotine Ecig-exposed groups also improved MCA dilation in the presence of TEMPOL compared to their respective ACh-only response (p<0.05, **Figure 4B**). Similar to Tempol, the addition of Febuxostat also improved MCA dilatory capacity to ACh but unlike Tempol the improvement was in a sex-dependent manner. Combined graphs with males and females are shown in **Figure 4C&D**. Male offspring (with or without nicotine) showed significant improvement in all Ecig exposed groups (restoring vascular function to control level responses) (p<0.05, **Supplement Figure 1A** no nicotine, **Supplement Figure 1C** nicotine). Female offspring only saw partial improvement, but only with 30W exposure (p<0.05, **Supplement Figure 1B** no nicotine, **Supplement Figure 1D** nicotine). No difference related to nicotine are observed. Myogenic MCA responses were also measured and found to not be different (**Supplement Figure 2A-D**).

**Figure 3:**
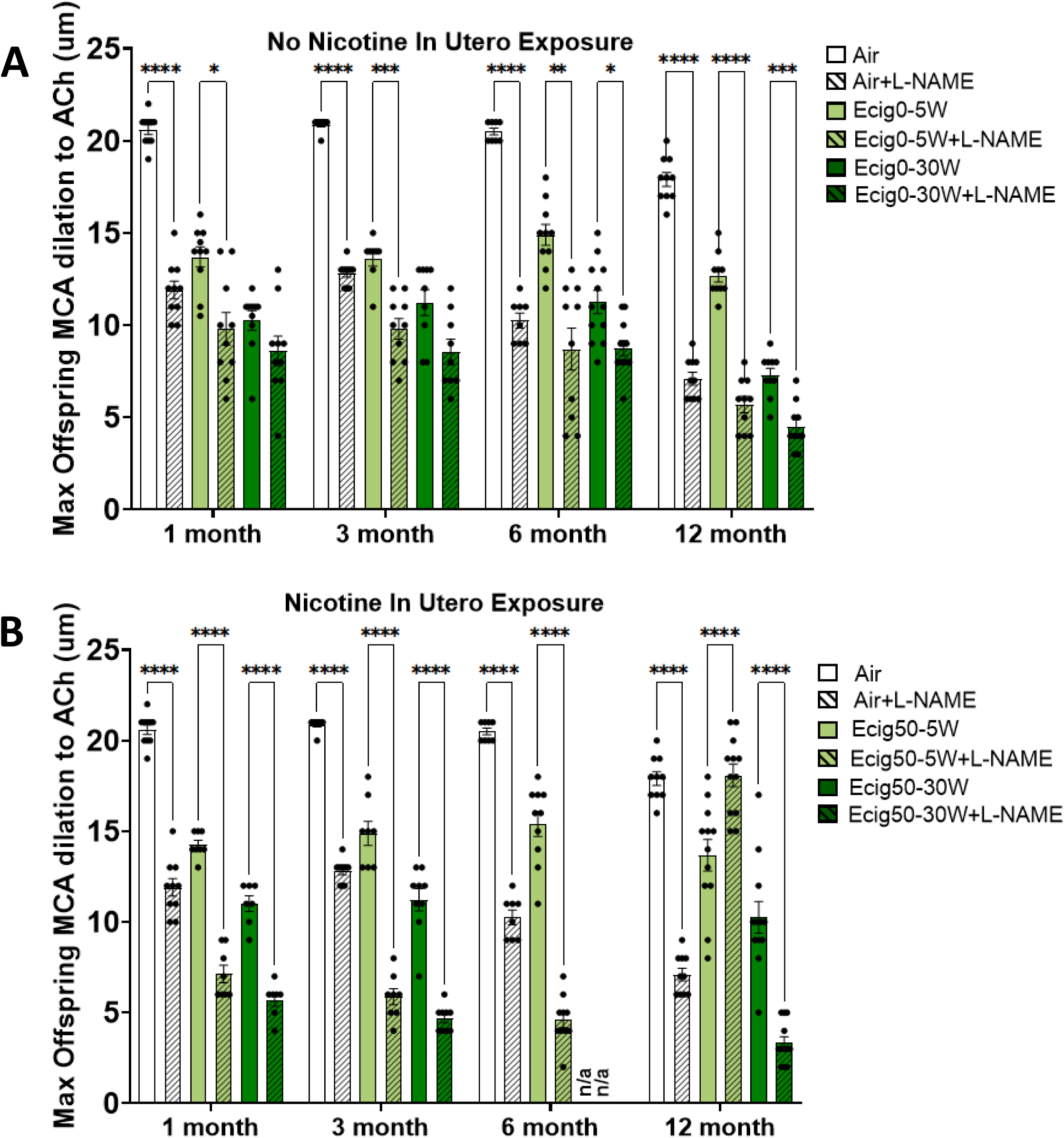
MCA response at 1-, 3-, 6-, 12-months to nitric oxide inhibitor (L-NAME) vasodilatory response was re-assessed with ACh separated by nicotine (**A**-No Nicotine, **B**-Nicotine) (n = 5- 6/group) Mean ± SE.

**Figure 4:**
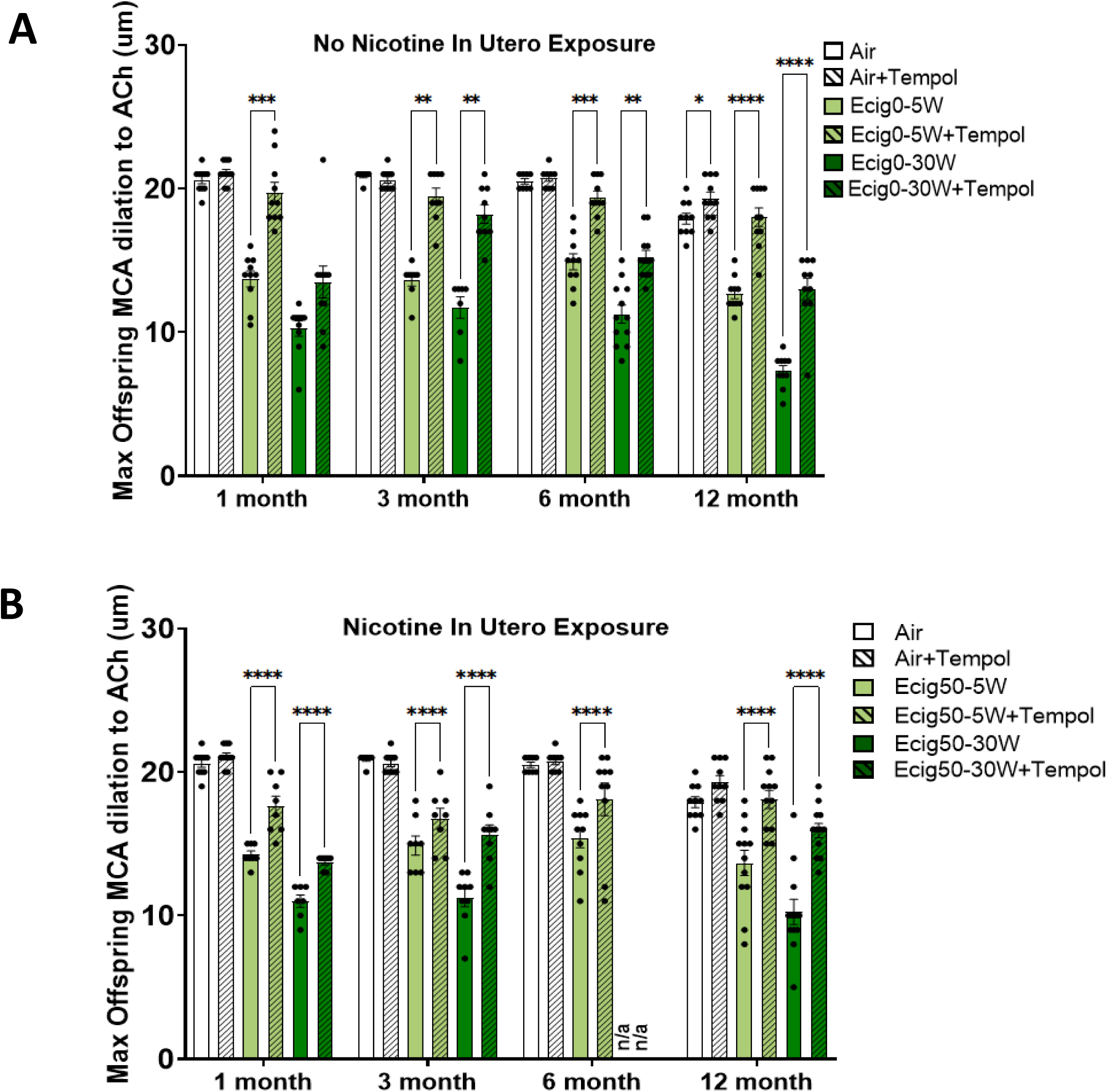
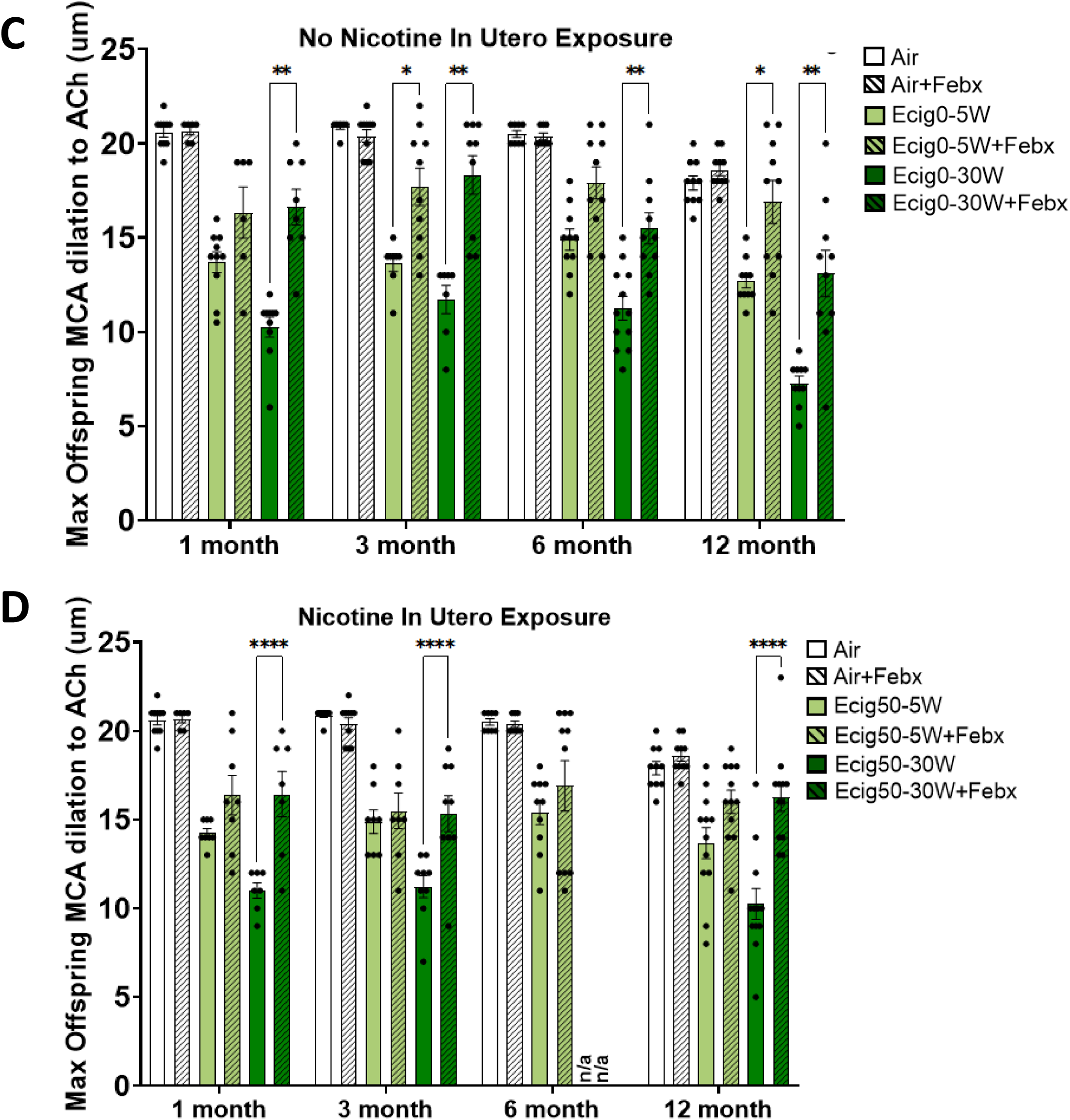
Offspring MCA at 1-, 3-, 6-, 12-months middle cerebral artery (MCA) maximal reactivity with incubation of a superoxide dismutase mimetic (TEMPOL) **A-**No nicotine**, B-**Nicotine, a xanthine oxidase inhibitor (Febuxostat) **C-**No nicotine**, D-**Nicotine, the vasodilatory response was re-assessed with ACh. (n = 5-6/group). Mean ± SE.

Neurocognitive and behavioral changes are reported in **Figures 5-7**, and **Supplemental Figure 3)**. Novel object recognition revealed offspring with maternal Ecig exposure to 30W (regardless of nicotine) led to short-term memory deficits as shown by a lack of preference for the novel object relative to the familiar (**Figure 5A&B**). Ecig50-5W group did not show a preference for the novel object. Barnes maze trials also indicate short-term memory deficits with a lack of improvement in path efficiency over the 5-day acquisition trial and a significant decrease in number of interactions with the escape hole during the reversal trial (p<0.05, **Figure 6A-C**). Lastly, Y-maze showed a decrease in percent alternation with a tendency to move around less (seen in heat map, **Figure 7A**) compared to Air (p<0.05, **Figure 7A&B**). Locomotor activity and anxiety-like behaviors were recorded at 3, 6, and 12 months using Open Field testing. We observed the Ecig offspring at 3-months, but not 6- and 12-months, showed decreased central tendency compared to Air (**Supplement Figure 3A&B**). However, at all ages, Ecig offspring had an increase in fecal boli compared to control (**Supplement Figure 3C**), suggesting greater level of anxiety compared to controls.

**Figure 5:**
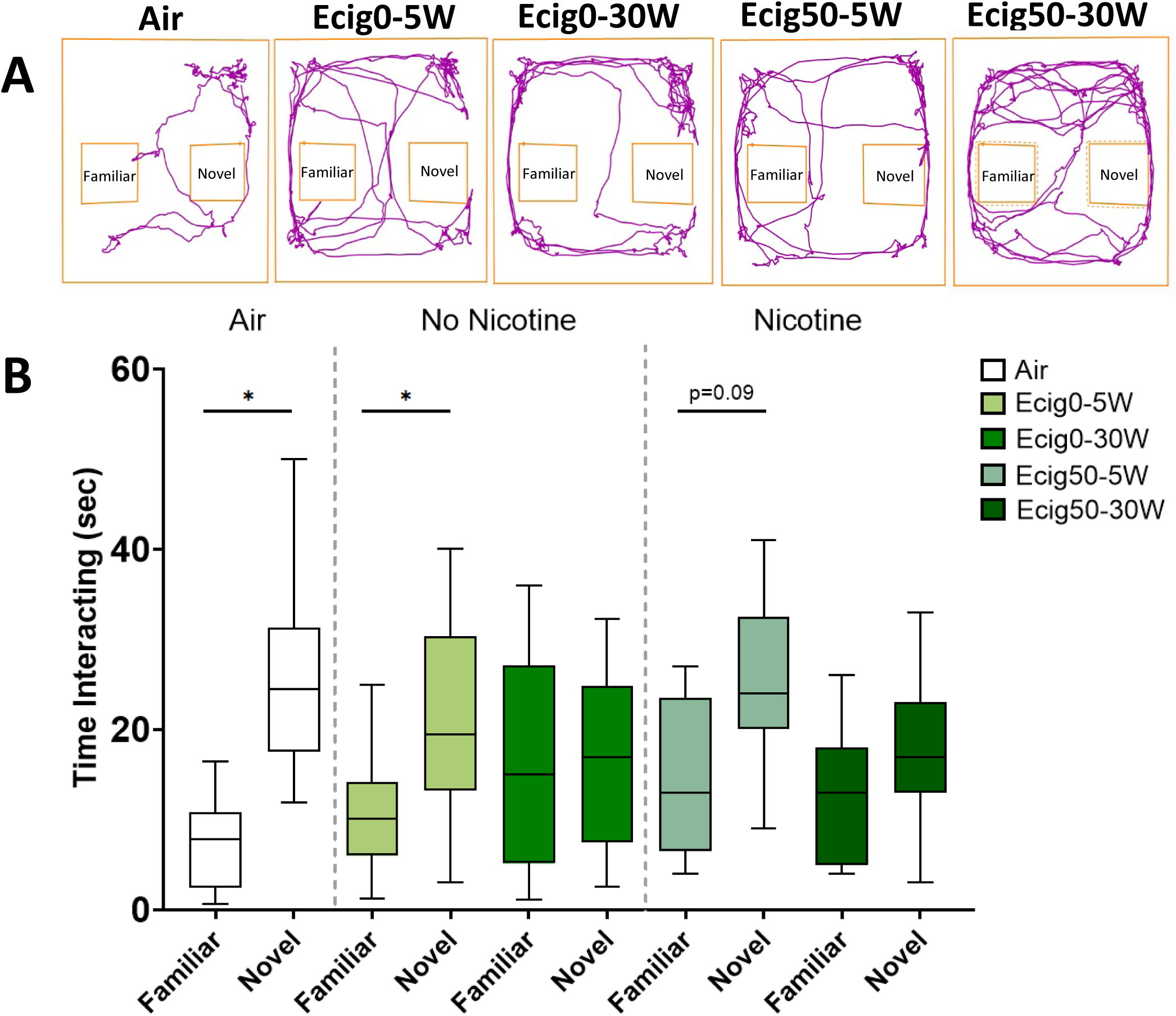
Novel object recognition in 12-month offspring movement track plots **A**, and number of interactions with familiar and novel object **B**. Mean ± SD.

**Figure 6:**
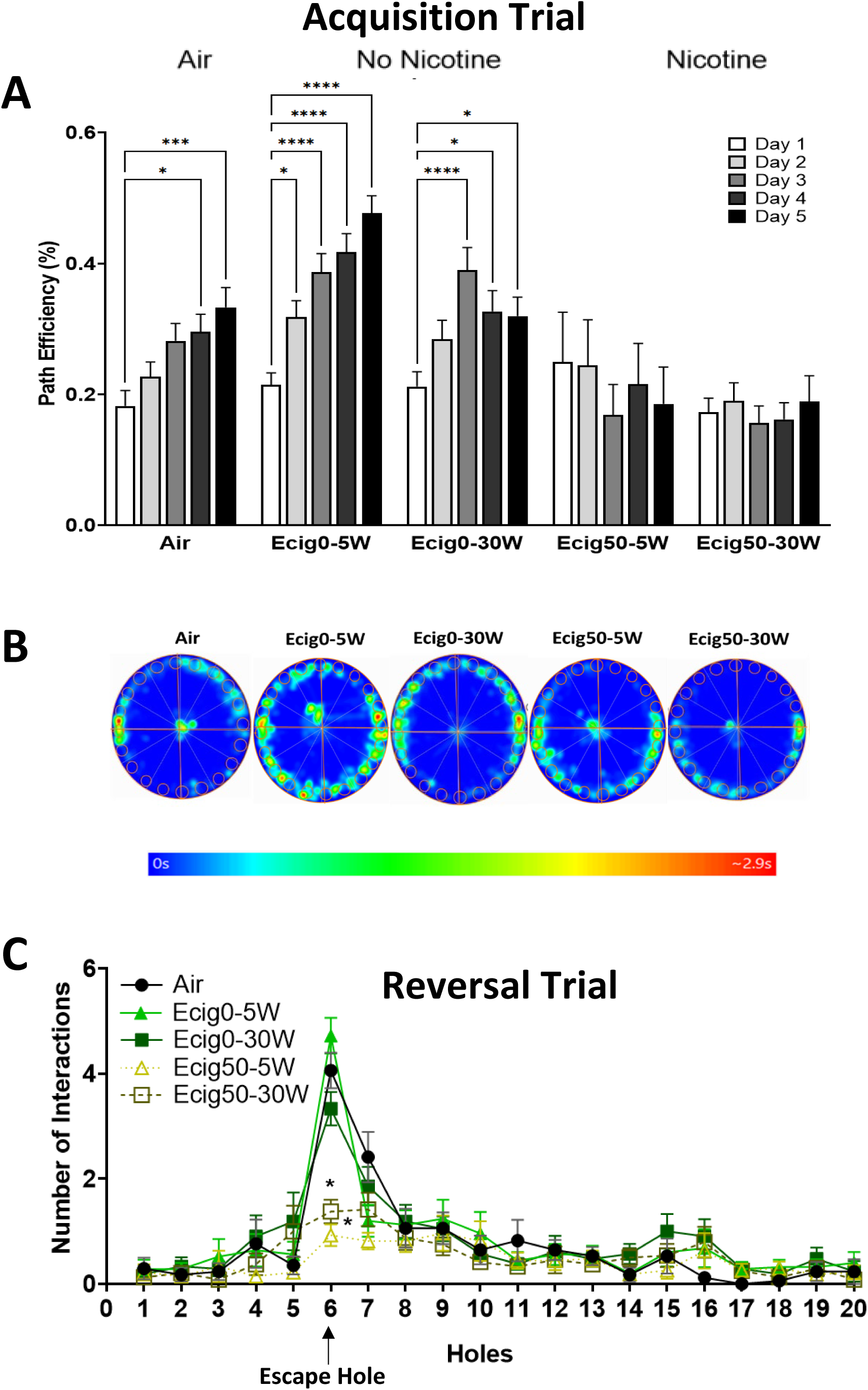
8-Month Barnes maze testing acquisition trial during 5-day training **A**, heat-maps showing time (sec) on the maze **B**, and number of interactions with each escape hole during reversal **C**. Mean ± SD.

**Figure 7:**
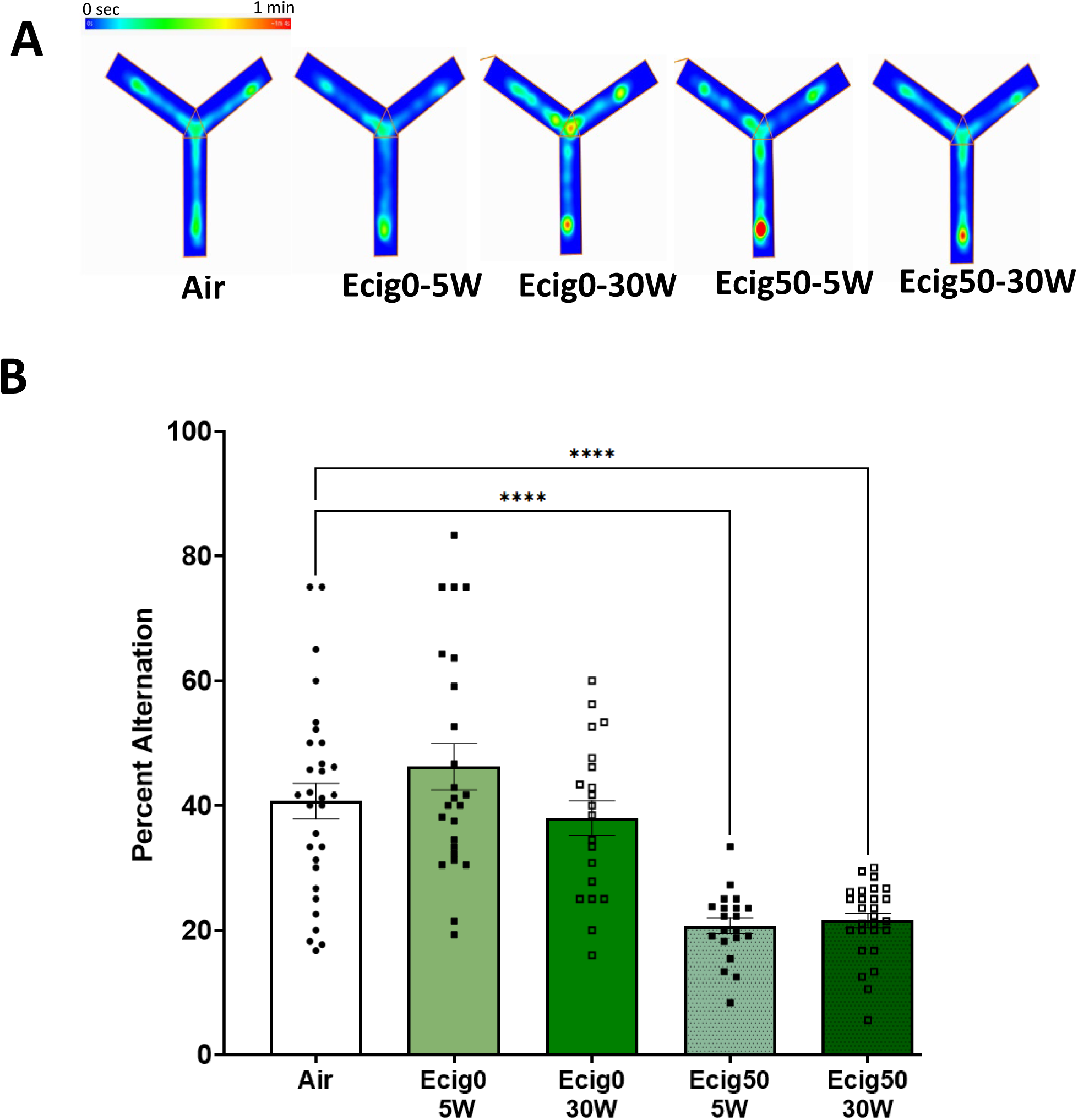
Heat-maps during Y-maze showing time (sec) on the maze **A**, percent alternation during Y-maze trial **B**. Mean ± SE.

EVs isolated from 12-month offspring plasma and characterized with the Nanosight analyses are shown in **Figure 8**. Here, offspring exposed to Ecig had a higher concentration of EV compared to Air (p<0.05, **Figure 8A&B**). There were no statistical differences between wattage or nicotine exposure.

**Figure 8:**
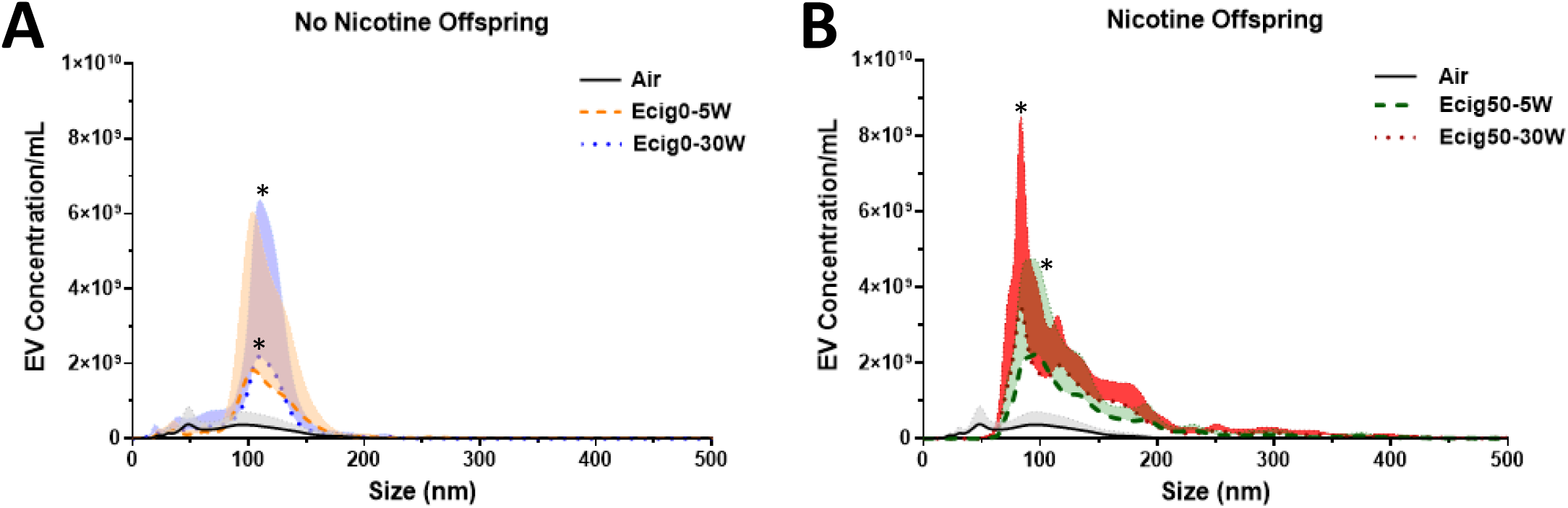
Extracellular Vesicles (EV) number and distribution from 12-month offspring plasma separated by nicotine (No nicotine **A**, Nicotine **B**) (n = 4-6/group). Mean ± SD.

Using ELISA-based assays, homogenate from the front brain revealed SIRT1 protein levels were significantly decreased in all Ecig groups compared to controls (p<0.05) at both 1- and 12-months of age (**Table 3**). At 12-months, we also found NOX1, Aꞵ1-40, and Aꞵ1-42 were increased in all Ecig groups (p<0.05, **Table 3**) except for Ecig50-5W. The ratio of Aꞵ1-42/Aꞵ1-40 was decreased in all Ecig-exposed offspring (regardless of wattage or nicotine exposure) (**Figure 9**). Correlations between NOX1 and SIRT1 had a R^2^ of 0.84 showing as SIRT1 decreases, NO1 levels increase (**Figure 9A**). Aꞵ1-40 and SIRT1 showed a similar correlation with a R^2^ of 0.79 (**Figure 9B**). Aꞵ1-42 did not show a strong correlation with SIRT1 levels R^2^=0.01 (**Figure 9C**). The ratio of Aꞵ1-42 to Aꞵ1-40 showed a significant decrease in all Ecig-exposed offspring (p<0.05, **Figure 9D**) except for Ecig50-30W (p=0.055).

**Figure 9:**
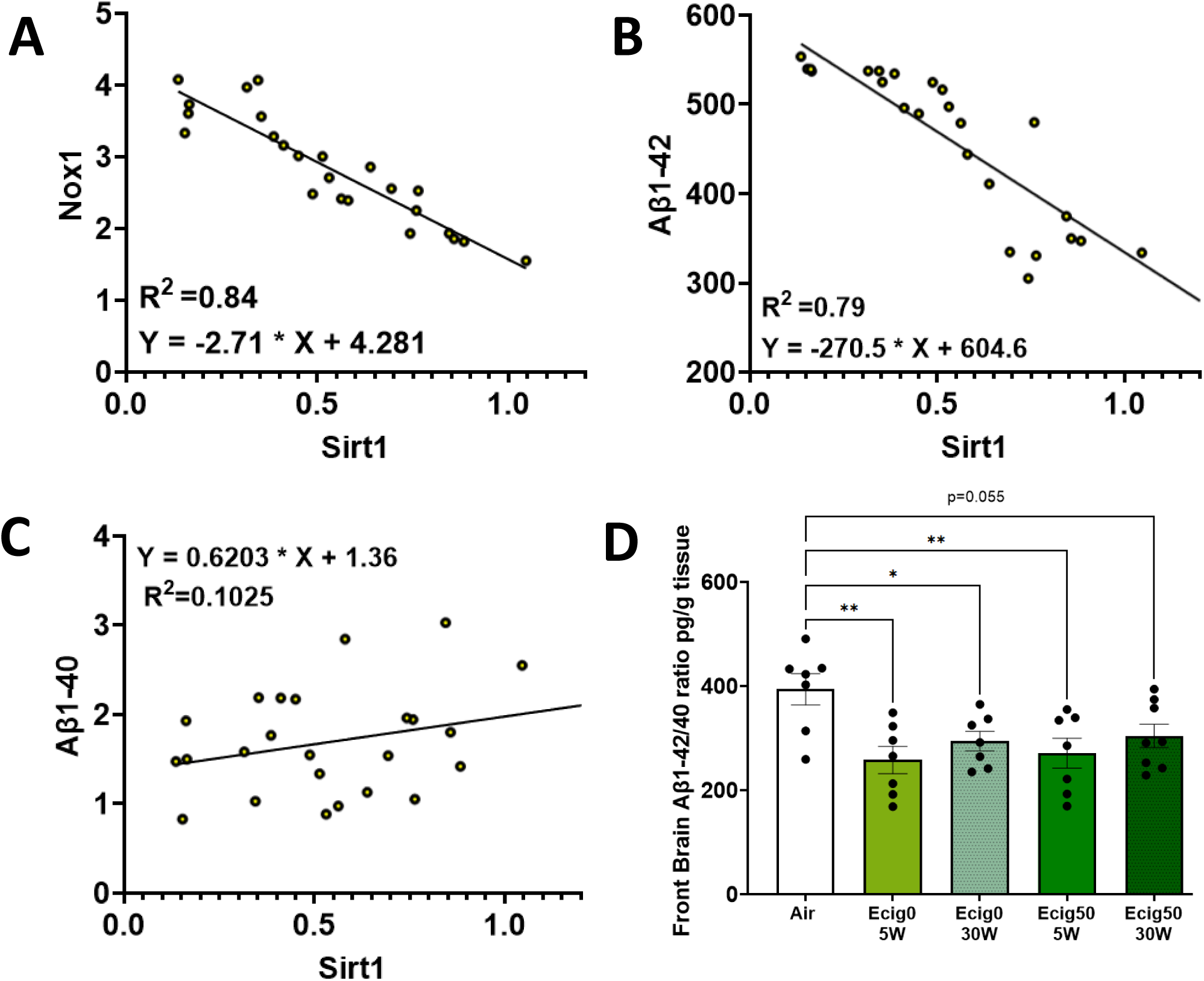
Sirt1 Relationship between NOX1 **A**, and Aꞵ1-42 **B**, Aꞵ1-40 **C**, Aꞵ1-42/40 Ratio **D** (n=8 offspring/group) Mean ± SD.

**Table 3:** Circulating and Neuronal Factors.

| **Offspring Circulating Factors** |  |  |  |  |  |
| --- | --- | --- | --- | --- | --- |
| Offspring n= | 8 | 8 | 8 | 8 | 8 |
| **1-Month** |  |  |  |  |  |
| Brain Tissue SIRT1<br>(ng/g) | 0.306 ± 0.06 | **0.153 ± 0.05\*** | **0.125 ± 0.03\*** | **0.158 ± 0.07\*** | **0.182 ± 0.06\*** |
| **12-Months** |  |  |  |  |  |
| Brain Tissue SIRT1<br>(ng/g) | 0.858 ± 0.26 | **0.427 ± 0.21\*** | **0.427 ± 0.15\*** | **0.475 ± 0.18\*** | **0.544 ± 0.22\*** |
| Brain Tissue NOX1<br>(ng/g) | 2.13 ± 0.49 | **3.01 ± 0.47\*** | 2.74 ± 0.67 | **3.02 ± 0.60\*** | **3.09 ± 0.78\*** |
| Brain Tissue Aβ1-40<br>(pg/g) | 1.10 ± 0.24 | **1.94 ± 0.60\*** | **1.86 ± 0.31\*** | **2.19 ± 0.62\*** | **1.79 ± 0.39\*** |
| Brain Tissue Aβ1-42<br>(pg/g) | 352 ± 60 | **461 ± 77\*** | **525 ± 9.5\*** | **501 ± 16\*** | **527 ± 34\*** |

Gene expression of ‘clock genes’ were measured using RT-PCR in brain homogenates of 1- and 12-month-old offspring (**Table 4**). Arnt1 was significantly increased in all Ecig groups compared to Air controls in 1-month old offspring (p<0.05, **Table 4**) but only in Ecig 50-5W in 12- month-old. The remaining genes (i.e. Per1, Per3, Clock, Cry1, and Cry2) were all increased with nicotine exposure (in both 5W and 30W) (p<0.05, **Table 4**). Clock and Cry1 were also increased Ecig0-30W (p<0.05), but not in Ecig0-5W (**Table 4**).

**Table 4:** Clock Genes.

|  | Air | Ecig0-5W | Ecig50-5W | Ecig0-30W | Ecig50-30W |
| --- | --- | --- | --- | --- | --- |
| Offspring n= | 5 | 5 | 5 | 5 | 5 |
| **1-Month** |  |  |  |  |  |
| Arnt1 | 0.10 ± 0.02 | **0.25 ± 0.13\*** | **0.27 ± 0.08\*** | **0.27 ± 0.05\*** | **0.29 ± 0.02\*** |
| Per1 | 0.16 ± 0.04 | 0.25 ± 0.09 | **0.32 ± 0.11\*** | 0.34 ± 0.12 | 0.30 ± 0.06 |
| Per3 | 0.36 ± 0.09 | 0.49 ± 0.19 | **0.51 ± 0.11\*** | 0.62 ± 0.13 | **0.71 ± 0.08\*** |
| Clock | 0.16 ± 0.02 | 0.26 ± 0.09 | **0.29 ± 0.09\*** | **0.31 ± 0.02\*** | **0.33 ± 0.04\*** |
| Cry1 | 0.13 ± 0.06 | 0.29 ± 0.12 | **0.36 ± 0.18\*** | **0.39 ± 0.06\*** | **0.34 ± 0.05\*** |
| Cry2 | 0.44 ± 0.05 | 0.66 ± 0.25 | **0.69 ± 0.24\*** | 0.86 ± 0.07 | **0.88 ± 0.27\*** |
| **12-Months** |  |  |  |  |  |
| Arnt1 | 0.23 ± 0.03 | 0.20 ± 0.05 | **0.20 ± 0.05\*** | 0.15 ± 0.01 | 0.20 ± 0.02 |
| Per1 | 0.30 ± 0.06 | 0.28 ± 0.07 | 0.33 ± 0.06 | 0.24 ± 0.07 | 0.34 ± 0.08 |
| Per3 | 0.47 ± 0.08 | 0.49 ± 0.15 | 0.53 ± 0.02 | 0.42 ± 0.10 | 0.43 ± 0.16 |
| Clock | 0.33 ± 0.05 | 0.34 ± 0.09 | 0.33 ± 0.05 | 0.29 ± 0.04 | 0.30 ± 0.04 |
| Cry1 | 0.23 ± 0.05 | 0.20 ± 0.05 | 0.21 ± 0.03 | 0.21 ± 0.02 | 0.24 ± 0.06 |
| Cry2 | 0.50 ± 0.08 | 0.51 ± 0.11 | 0.53 ± 0.04 | 0.51 ± 0.11 | 0.45 ± 0.11 |

Coronal sections (with left and right hemispheres) from 12-month-old offspring showed an increase in silver stain in the cortex and total silver stain in nicotine-exposed offspring compared to Air controls (p<0.05, **Figure 10A&B**). In addition to neuronal damage, astrocyte, and endothelial cell co-staining showed an increase in astrocyte number in all Ecig groups compared to controls (p<0.05, **Figure 10A**), and the number of interactions between astrocytes and endothelial cells was increased in the 30W groups with and without nicotine compared to controls (p<0.05, **Figure 10B**).

**Figure 10:**
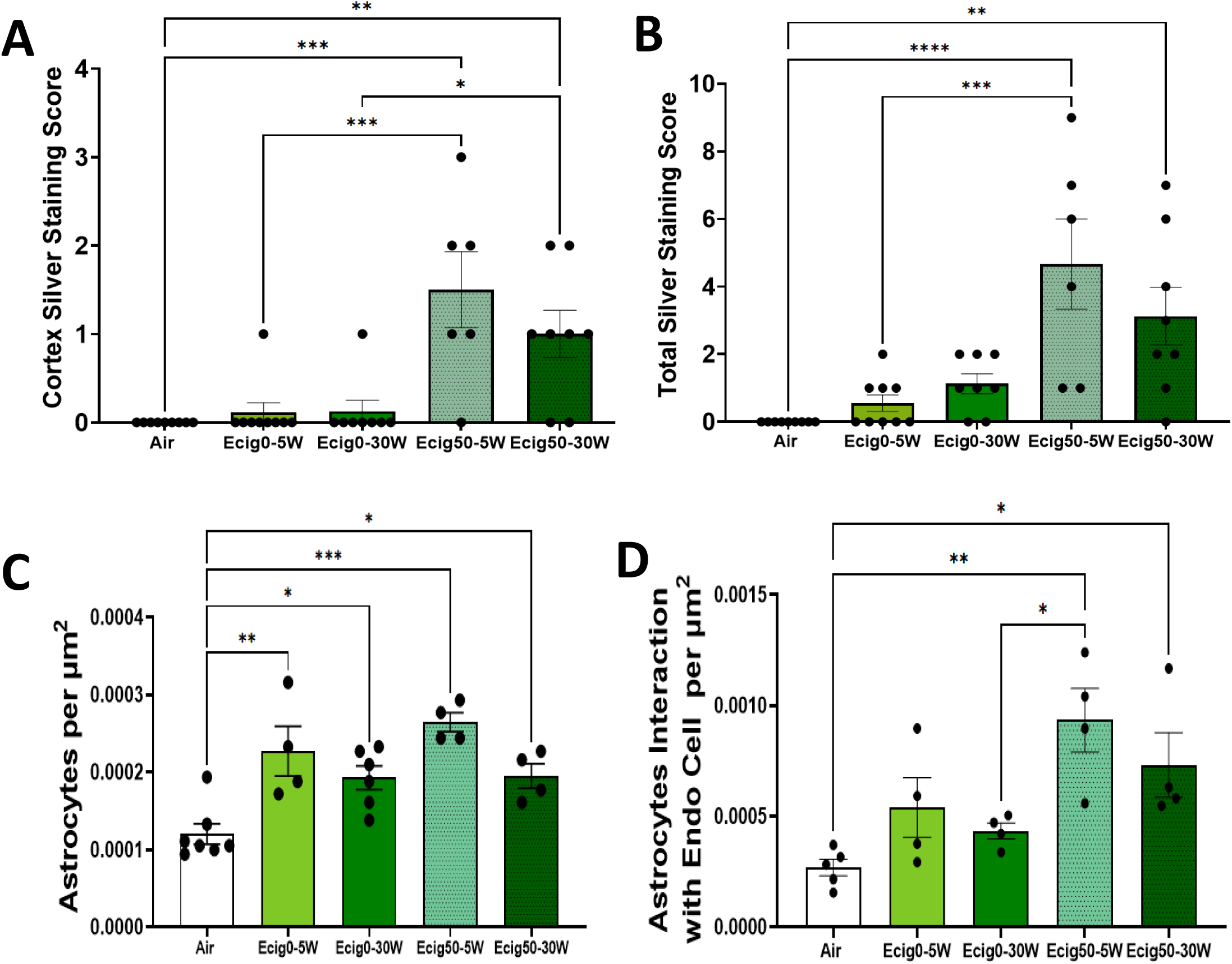
Coronal section scores from the cortex **A**, and Total brain **B**, number of astrocytes **C**, and number of interactions of astrocytes with endothelial cells **D**, from 12-month offspring coronal sections (n = 6-11 offspring/group). Mean ± SE.

## DISCUSSION

The main findings from this study are, 1) maternal vaping, with or without nicotine, results in long-term vascular dysfunction that persists in the adult life of offspring, 2) Ecig aerosol created from either 5W and 30W both results in significant endothelial cell dysfunction, but the effect is worse with higher wattage, 3) nicotine does not appear to be involved in the etiology of vascular dysfunction, but maternal vaping with nicotine resulted in significant neurocognitive and memory impairments in young and adult offspring, and 4) in utero nicotine exposure from Ecigs alters gene and protein expression in the brain in a manner consistent with accelerated cellular senescence and increases risk/biomarkers for neurogenerative disease compared controls.

As previously reported [22, 23] this study found offspring with maternal exposure to Ecigs had vascular impairment and the severity of impairment was dependent on device setting (wattage) rather than nicotine. However, to our knowledge, this is the first study to follow and report offspring up to 12-months of age – and thus show this effect persists into mid-life (and likely beyond). In addition to cerebrovascular deficits, offspring had memory and neuronal damage leading to the activation of astrocytes and increased interaction between astrocytes and endothelial cells. Exposure to nicotine has been shown to impact neuronal health and activate glial cells [49]. In our study, nicotine seemed to be the driver for neuronal damage with only the nicotine groups (regardless of the wattage) having more damage in the cortex and overall brain. While nicotine *per se* did not appear to correlate with the increase in astrocytes, the nicotine-exposed group did have an increase in the number of astrocytes interacting with endothelial cells. It remains unclear if this is an attempt to aid the integrity of the BBB, or a maladaptive response of the neurovascular network in the brain.

Neuroinflammation can occur with mitochondrial dysfunction, DNA damage, and/or with elevated ꞵ-amyloid [50, 51]. Astrocytes help to protect the brain by removing the source of the inflammatory response and repairing affected tissue, but with chronic inflammation, the reverse can happen. Recent studies have shown that astrocytes express nicotinic acetylcholine receptors (nAChRs) and mediate calcium signaling [52]. Exposure to nicotine can cause a rapid increase in the number of processes and eventually an increase in cell bodies [52]. Others find exposure to cigarette smoke leads to changes to the astrocytes of the hippocampus and astrocytes when exposed to a secondary insult (such as a traumatic brain injury) are further activated compared to cigarette smoke alone [53]. These astrocyte changes induced by cigarette smoke in combination with a traumatic brain injury may contribute to poorer cognitive outcomes and in particular memory deficits [54]. Environmental (e.g. cigarette smoke) and endogenous sources (e.g. epigenetic changes) can lead to chronic inflammation and activate astrocytes [55, 56]. If these glial cells remain activated this can lead to neurodegenerative diseases [51], such as Alzheimer’s disease and Parkinson’s disease. Activation of astrocytes occurs via the nuclear factor kappa-light-chain-enhancer of activated B cells (NF-κB) pathway [57]. Both cigarettes [58] and Ecig [59] have been shown to activate NF-κB. Further, NF-κB has also been linked to SIRT1, for example, NF-κB can down-regulate SIRT1 through the expression of miR-34a, IFNγ, and reactive oxygen species [60]. Production of ROS and other factors in turn stimulate NF-κB creating an inflammatory cycle and the sets the stage for age-related diseases [61]. NF-κB is driving a pro-inflammatory phenotype with glycolytic metabolism, whereas SIRT1 supports oxidative respiration and anti-inflammatory responses [62, 63]. However, activation of SIRT1 also inhibits NF-κB through direct deacetylation of p65 subunit and indirectly through activation of PPARα [60]. Thus, it is tempting to speculate NF-kB could be a potential mechanism for the activation of astrocytes in our study. It is interesting to note that our data from Ecigs are consistent with prior work reporting that maternal tobacco smoke exposure alters fetal programming and that offspring have an increase in neuroinflammatory response leading to activation of primary astrocytes [64].

Our offspring had decreased SIRT1 protein concentration at both 1- and 12-months of age. SIRT1 is an NAD+ dependent deacetylase important for the maintenance of histone acetylation, and prominent in the activation of gene expression and epigenetic regulation. Loss of SIRT1 whether due to an insult such as Ecig exposure or a genetic knockout out has been shown to impact behavior and circadian rhythm [65]. Offspring demonstrated both neurocognitive deficits and alterations in clock genes. Further, SIRT1 mediates the metabolism of energy sources in metabolically active tissues, [66, 67] and can interact with insulin signaling pathways, and activation of SIRT1 is linked with an increase in insulin sensitivity. When offspring weight was recorded, initial measurements (at weaning) showed offspring were significantly smaller in lean and fat mass compared to controls, but as they age the Ecig offspring surpassed controls suggesting a metabolic dysregulation commonly seen with fetal growth restriction. SIRT1 can promote the mobilization of fat from white adipose tissue [68], therefore less SIRT1 can mean retention of more fat tissue and thus greater body mass. If there is an NAD+ deficiency then SIRT function is affected, therefore it is thought to be a functional link between metabolic activity and genome stability as well as aging [69]. It is notable that NF-κB can mediate the expression of the NOX family. NF-κB can aggravate TNF-α by regulating the oxidative stress response and the expression NOX, thereby NF-κB could promote the NOX1 transcriptional activity via binding its promoter [70]. NOX is an enzyme complex that is a primary source of ROS generation [71, 72]. It is important to recognize that not all ROS is bad as it can be used for cellular signaling. Still an imbalance can result in oxidative stress which has been implicated in cardiovascular disease, neurodegeneration, and aging. In addition to a decrease in SIRT1, offspring had a general increase in NOX1 in the brain, this combined with the rescue to vascular function when vessels are treated with Tempol or Febuxostat suggests that ROS is certainly a driver of endothelial dysfunction. SIRT1 is known to be anti-inflammatory and viewed as cardioprotective with its regulation of eNOS [73, 74], thus connecting it to NO bioavailability. The fact that we find significant correlations between SIRT1, NOX1, and ꞵ-amyloid suggests these factors are likely interrelated, but further work is needed to fully understand these relationships in association with vaping.

There is emerging evidence that SIRT1 is also implicated in Alzheimer’s progression because of its known neuroprotective ability to promote neuronal survival, regulation of ꞵ-amyloid metabolism, and modulation of tau phosphorylation – all of which are important for Alzheimer’s pathology [75]. Exposure to cigarette smoke is known to increase the risk for the development of Alzheimer’s disease [76], but with cigarettes, it is difficult (if not impossible) to separate inhalation of nicotine from other constituent in the smoke. In our study, we see that most of our behavior tests (i.e. Barnes and Y-maze) show that cognitive declines are nicotine-mediated. Our findings are consistent with a wide array of literature (mostly from nicotine delivered by dermal or injections) prenatal or perinatal exposure upregulates nAChRs and thus alters cell proliferation, differentiation, and myelin formation during embryo and fetus development [77–80]. This change alters the brain circuitry, response to neurotransmitters, and even brain volume. Specifically, exposure to nicotine during development reduces neurogenesis in the hippocampus, impacting learning/memory [81] and sensory cortical processing as well as reward processing in the nucleus accumbens. These changes in the circuitry are preserved in both rats and humans [27], and when human data reports prenatal nicotine exposure, offspring have an increased risk for adverse neurodevelopmental disorders such as Attention-Deficit Hyperactivity (ADHD), anxiety, and depression. Our findings suggest that in utero exposure to Ecigs containing nicotine are likely to have same adverse neurocognitive effects.

## Conclusions

In this study we report the role nicotine plays with maternal Ecig exposure on adult offspring vascular and neurocognitive function. While the responses in these data are nearly identical between males and females, we did observe a sex-dependent responses when blood vessels were treated with Febuoxstat, suggesting estrogen may have negative influence with the xanthine oxidase pathway. Nonetheless, consistent with our previous studies we find that the principal chemical of concern when evaluating vascular endothelial dysfunction in not nicotine. Importantly, we observe that nicotine did influence and impact behavior and severity of neuronal damage, demonstrating that although might not greatly influence vascular reactivity it does have (and widely known in the literature) to have significant adverse effects in offspring neurodevelopment. Regardless of wattage or nicotine, offspring in the Ecig groups had evidence of a decrease in SIRT1 and elevated oxidative stress (i.e. NOX1). Interestingly, offspring exhibit Alzheimer’s-like pathology with an increase in ꞵ-amyloid and an overall decrease in Aꞵ1-42:40 ratio. Evidence for neuroinflammation (increased cerebral silver staining, increased astrocytes) were also seen in offspring, suggest that there is significant harm stemming solely from the base solution. The overall phenotype observed from in offspring with in-utero exposure to Ecigs appears to be consistent with advanced cellular and vascular aging when compared to controls. These pre-clinical data provide early evidence that maternal vaping during pregnancy is not safe for the long-term vascular and neurocognitive health of offspring.

## Author Contributions

AM and IMO conceived and designed the study; AM and DC conducted experiments, collected data, and/or performed daily exposures; AM and IMO analyzed data and interpreted results of the experiment; AM, PDC, and IMO prepared figures and drafted the manuscript; all authors help to edit, revise, and approve the final version of the manuscript.

---

**Supplement Figure 1:**
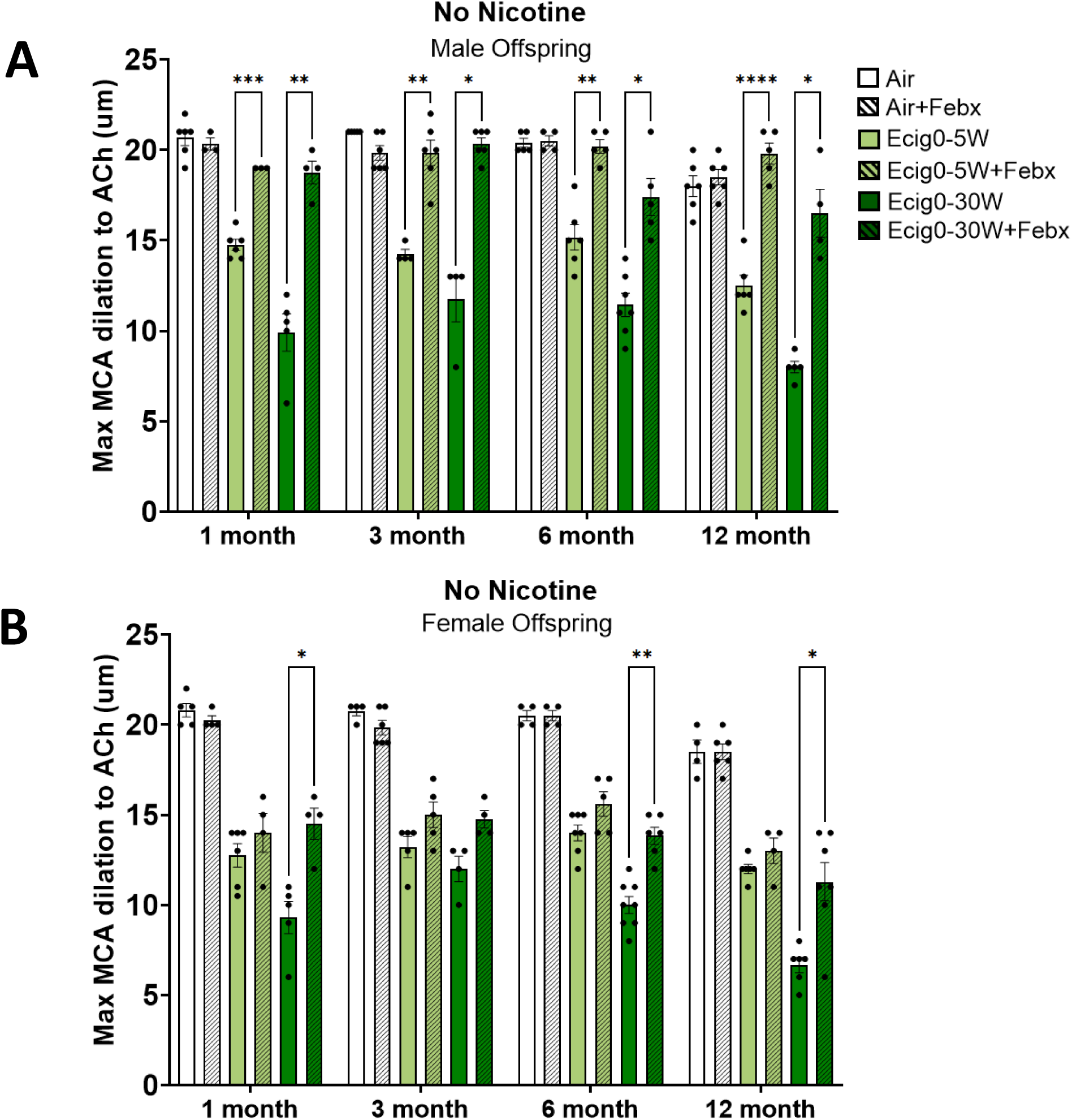
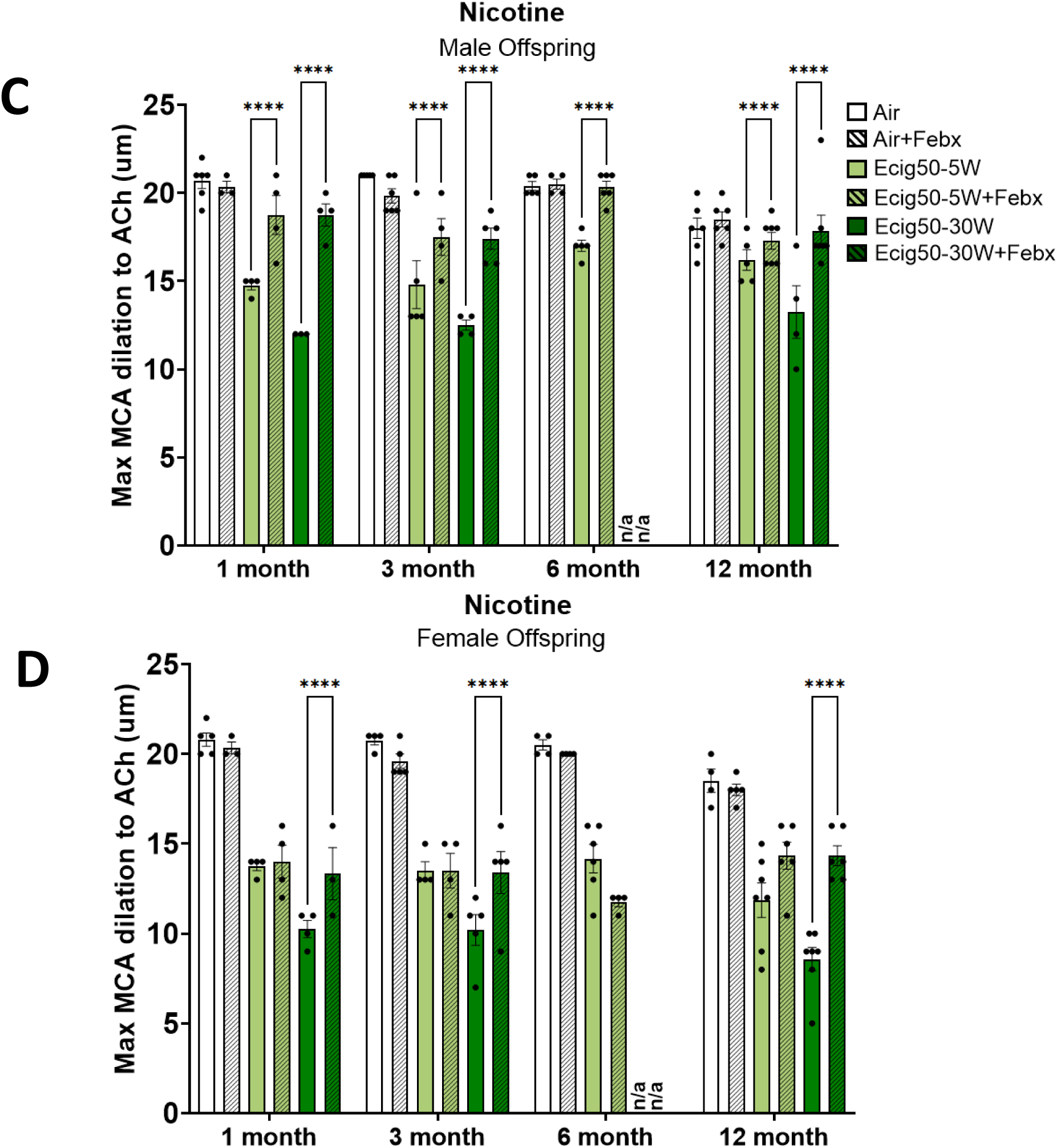
Febuxostat sex dependent response. Male offspring **A-**No Nicotine &**C-** Nicotine, Female offspring **B-**No nicotine **D-**Nicotine, the vasodilatory response was re-assessed with ACh. (n = 5-6/group). Mean ± SE.

**Supplement Figure 2:**
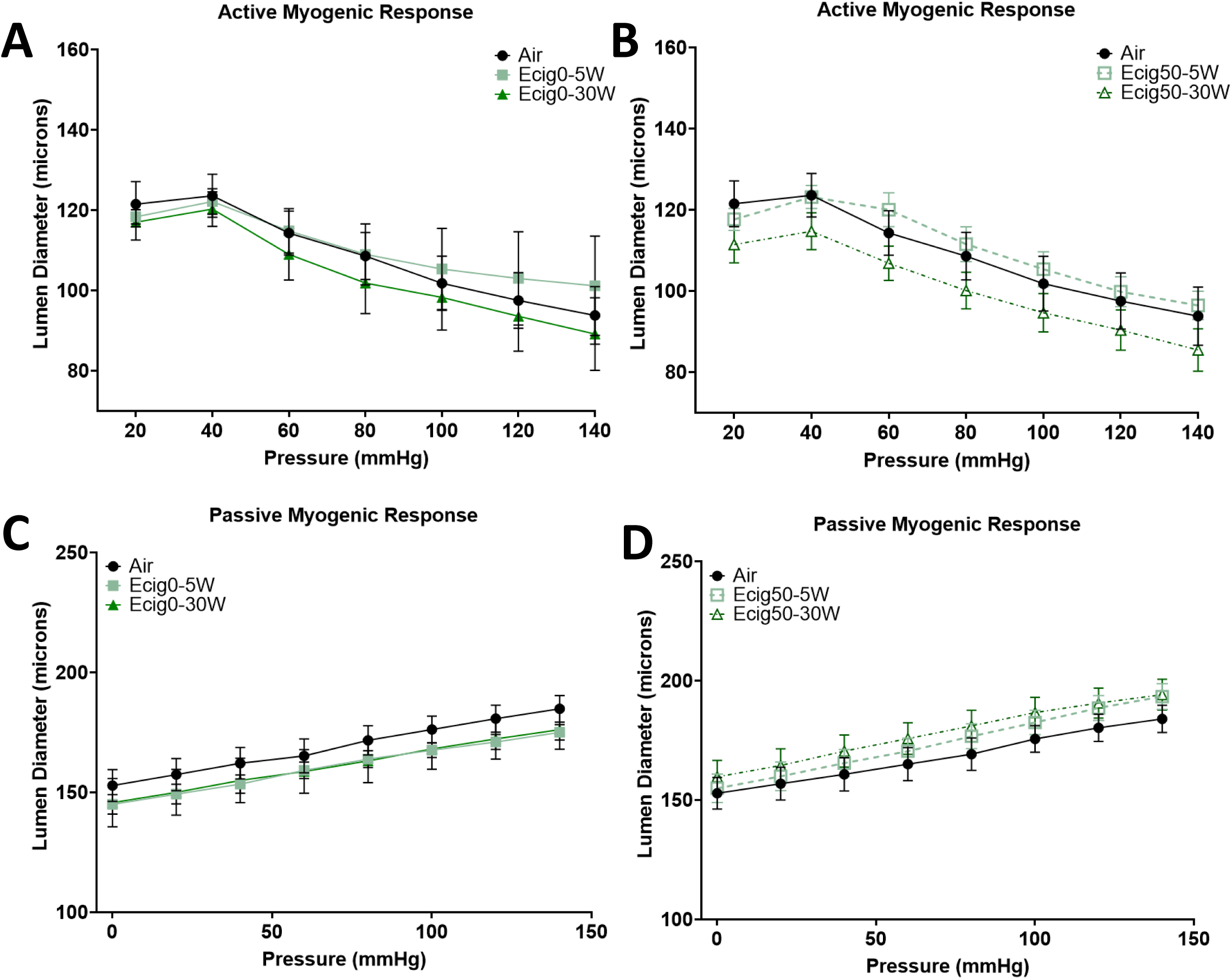
Active **A-**No nicotine**, B-**Nicotine and passive **C-**No nicotine**, D-**Nicotine myogenic response in 12-month offspring following MCA reactivity measurements separated by nicotine. (n = 5-6/group) Mean ± SD.

**Supplement Figure 3:**
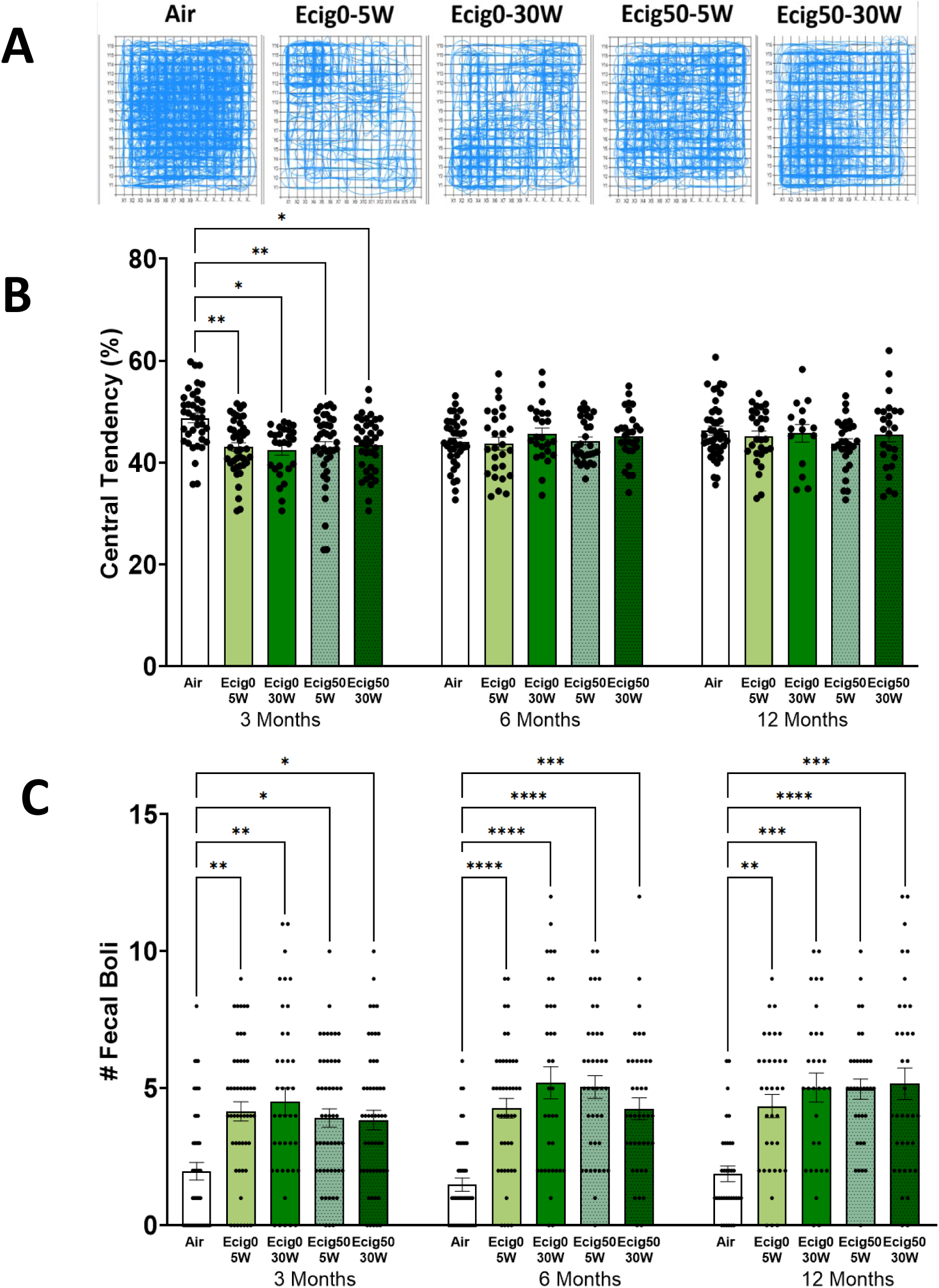
Anxiety-like behavior assessed with open field track plots **A**, central tendency **B**, Fecal Boli **C**. Mean ± SE.

## REFERENCES

1. Williams, M., et al., Metal and silicate particles including nanoparticles are present in electronic cigarette cartomizer fluid and aerosol. PLoS One, 2013. 8(3): p. e57987.

2. Olmedo, P., et al., Metal Concentrations in e-Cigarette Liquid and Aerosol Samples: The Contribution of Metallic Coils. Environ Health Perspect, 2018. 126(2): p. 027010.

3. Mikheev, V.B., et al., Real-Time Measurement of Electronic Cigarette Aerosol Size Distribution and Metals Content Analysis. Nicotine Tob Res, 2016. 18(9): p. 1895–902.

4. Conklin, D.J., et al., Electronic cigarette-generated aldehydes: The contribution of e-liquid components to their formation and the use of urinary aldehyde metabolites as biomarkers of exposure. Aerosol Sci Technol, 2018. 52(11): p. 1219–1232.

5. El-Hellani, A., et al., Nicotine and Carbonyl Emissions From Popular Electronic Cigarette Products: Correlation to Liquid Composition and Design Characteristics. Nicotine Tob Res, 2018. 20(2): p. 215–223.

6. Jensen, R.P., et al., Hidden Formaldehyde in E-Cigarette Aerosols. New England Journal of Medicine, 2015. 372(4): p. 392–394.

7. Kosmider, L., et al., Carbonyl Compounds in Electronic Cigarette Vapors: Effects of Nicotine Solvent and Battery Output Voltage. Nicotine & Tobacco Research, 2014. 16(10): p. 1319–1326.

8. Hansen, J.M., Oxidative stress as a mechanism of teratogenesis. Birth Defects Res C Embryo Today, 2006. 78(4): p. 293–307.

9. Dennery, P.A., Effects of oxidative stress on embryonic development. Birth Defects Res C Embryo Today, 2007. 81(3): p. 155–62.

10. Greene, R.M. and M.M. Pisano, Developmental toxicity of e-cigarette aerosols. Birth Defects Research, 2019. 111(17): p. 1294–1301.

11. Khan, N.A., et al., Waterpipe smoke and e-cigarette vapor differentially affect circadian molecular clock gene expression in mouse lungs. PLoS One, 2019. 14(2): p. e0211645.

12. Yoon, C., C.P. May, and L. Hasher, Aging, circadian arousal patterns, and cognition, in Cognition, aging and self-reports. 1998, Psychology Press. p. 113–136.

13. Orzabal, M.R., et al., Chronic exposure to e-cig aerosols during early development causes vascular dysfunction and offspring growth deficits. Transl Res, 2019. 207: p. 70–82.

14. Mills, A., et al., Short-term effects of electronic cigarettes on cerebrovascular function: A time course study. Exp Physiol, 2022. 107(8): p. 994–1006.

15. Mills, A., et al., Effects of electronic cigarette E-liquid and device wattage on vascular function. Toxicol Appl Pharmacol, 2023. 474: p. 116631.

16. Rahman, M.A., et al., E-cigarettes and smoking cessation: evidence from a systematic review and meta-analysis. PLoS One, 2015. 10(3): p. e0122544.

17. Rose, J.J., et al., Cardiopulmonary Impact of Electronic Cigarettes and Vaping Products: A Scientific Statement From the American Heart Association. Circulation, 2023. 148(8): p. 703–728.

18. Morris, P.B., et al., Cardiovascular Effects of Exposure to Cigarette Smoke and Electronic Cigarettes: Clinical Perspectives From the Prevention of Cardiovascular Disease Section Leadership Council and Early Career Councils of the American College of Cardiology. J Am Coll Cardiol, 2015. 66(12): p. 1378–91.

19. Buchanan, N.D., et al., Cardiovascular risk of electronic cigarettes: a review of preclinical and clinical studies. Cardiovasc Res, 2020. 116(1): p. 40–50.

20. Tarran, R., et al., *E-Cigarettes and Cardiopulmonary Health.* Function, 2021. 2(2).

21. Neczypor, E.W., et al., E-Cigarettes and Cardiopulmonary Health: Review for Clinicians. Circulation, 2022. 145(3): p. 219–232.

22. Aboaziza, E., et al., Maternal electronic cigarette use during pregnancy affects long-term arterial function in offspring. J Appl Physiol (1985), 2023. 134(1): p. 59–71.

23. Burrage, E.N., et al., Long-term cerebrovascular dysfunction in the offspring from maternal electronic cigarette use during pregnancy. Am J Physiol Heart Circ Physiol, 2021. 321(2): p. H339–H352.

24. Jones, K. and G.A. Salzman, The Vaping Epidemic in Adolescents. Mo Med, 2020. 117(1): p. 56–58.

25. Dahlin, S., et al., Maternal tobacco use and extremely premature birth - a population-based cohort study. BJOG, 2016. 123(12): p. 1938–1946.

26. Al-Sawalha, N., et al., Effect of electronic cigarette aerosol exposure during gestation and lactation on learning and memory of adult male offspring rats. Physiol Behav, 2020. 221: p. 112911.

27. Liu, F., et al., Maternal Nicotine Exposure During Gestation and Lactation Period Affects Behavior and Hippocampal Neurogenesis in Mouse Offspring. Front Pharmacol, 2019. 10: p. 1569.

28. Navarro, H.A., et al., Prenatal exposure to nicotine impairs nervous system development at a dose which does not affect viability or growth. Brain Res Bull, 1989. 23(3): p. 187–92.

29. Ozekin, Y.H., et al., Intrauterine exposure to nicotine through maternal vaping disrupts embryonic lung and skeletal development via the potassium channel. Developmental Biology, 2023. 501: p. 111–123.

30. Olfert, I.M., et al., Chronic exposure to electronic cigarettes results in impaired cardiovascular function in mice. J Appl Physiol (1985), 2018. 124(3): p. 573–582.

31. Mohammadi, L., et al., Chronic E-Cigarette Use Impairs Endothelial Function on the Physiological and Cellular Levels. Arterioscler Thromb Vasc Biol, 2022. 42(11): p. 1333–1350.

32. Rao, P., et al., Comparable Impairment of Vascular Endothelial Function by a Wide Range of Electronic Nicotine Delivery Devices. Nicotine & Tobacco Research, 2022. 24(7): p. 1055–1062.

33. Rao, P., J. Liu, and M.L. Springer, JUUL and Combusted Cigarettes Comparably Impair Endothelial Function. Tobacco Regulatory Science, 2020. 6(1): p. 30–37.

34. Benowitz, N.L. and A.D. Burbank, Cardiovascular toxicity of nicotine: Implications for electronic cigarette use. Trends Cardiovasc Med, 2016. 26(6): p. 515–23.

35. O’Hara, B.F., et al., Nicotine and nicotinic receptors in the circadian system. Psychoneuroendocrinology, 1998. 23(2): p. 161–73.

36. Stevens, R.G., et al., Breast cancer and circadian disruption from electric lighting in the modern world. CA Cancer J Clin, 2014. 64(3): p. 207–18.

37. Yan, J., et al., Analysis of gene regulatory networks in the mammalian circadian rhythm. PLoS Comput Biol, 2008. 4(10): p. e1000193.

38. Robinson, I. and A.B. Reddy, Molecular mechanisms of the circadian clockwork in mammals. FEBS Lett, 2014. 588(15): p. 2477–83.

39. Hirayama, J., et al., CLOCK-mediated acetylation of BMAL1 controls circadian function. Nature, 2007. 450(7172): p. 1086–90.

40. Lechasseur, A., et al., Exposure to electronic cigarette vapors affects pulmonary and systemic expression of circadian molecular clock genes. Physiol Rep, 2017. 5(19).

41. Hwang, J.W., et al., Circadian clock function is disrupted by environmental tobacco/cigarette smoke, leading to lung inflammation and injury via a SIRT1-BMAL1 pathway. Faseb j, 2014. 28(1): p. 176–94.

42. Yao, H., et al., Disruption of Sirtuin 1-Mediated Control of Circadian Molecular Clock and Inflammation in Chronic Obstructive Pulmonary Disease. Am J Respir Cell Mol Biol, 2015. 53(6): p. 782–92.

43. Ding, X., et al., SIRT1 is a regulator of autophagy: Implications for the progression and treatment of myocardial ischemia-reperfusion. Pharmacol Res, 2024. 199: p. 106957.

44. Csiszar, A., et al., Oxidative stress and accelerated vascular aging: implications for cigarette smoking. Front Biosci (Landmark Ed), 2009. 14(8): p. 3128–44.

45. Grundy, D., Principles and standards for reporting animal experiments in The Journal of Physiology and Experimental Physiology. Exp Physiol, 2015. 100(7): p. 755–8.

46. Shao, X.M., et al., A mouse model for chronic intermittent electronic cigarette exposure exhibits nicotine pharmacokinetics resembling human vapers. J Neurosci Methods, 2019. 326: p. 108376.

47. Baumbach, G.L. and M.A. Hajdu, Mechanics and composition of cerebral arterioles in renal and spontaneously hypertensive rats. Hypertension, 1993. 21(6 Pt 1): p. 816–26.

48. Hughes, R.N., The value of spontaneous alternation behavior (SAB) as a test of retention in pharmacological investigations of memory. Neurosci Biobehav Rev, 2004. 28(5): p. 497–505.

49. Othman, M.A., A.M. Oseily, and E.M. Ramadan, Effects of Nicotine Administration on the Structure of Auditory Cortex of Adolescent Male Guinea Pigs, a Histological and Ultrastructural Study. Egyptian Journal of Histology, 2020. 43(3): p. 808–818.

50. Jellinger, K.A., Basic mechanisms of neurodegeneration: a critical update. J Cell Mol Med, 2010. 14(3): p. 457–87.

51. Wyss-Coray, T. and L. Mucke, Inflammation in neurodegenerative disease--a double-edged sword. Neuron, 2002. 35(3): p. 419–32.

52. Aryal, S.P., et al., Nicotine induces morphological and functional changes in astrocytes via nicotinic receptor activity. Glia, 2021. 69(8): p. 2037–2053.

53. Dobric, A., et al., Cigarette Smoke Exposure Induces Neurocognitive Impairments and Neuropathological Changes in the Hippocampus. Front Mol Neurosci, 2022. 15: p. 893083.

54. Ratliff, W.A., et al., Sidestream Smoke Affects Dendritic Complexity and Astrocytes After Model Mild Closed Head Traumatic Brain Injury. Cell Mol Neurobiol, 2022. 42(5): p. 1453–1463.

55. Stephenson, J., et al., Inflammation in CNS neurodegenerative diseases. Immunology, 2018. 154(2): p. 204–219.

56. Glass, C.K., et al., Mechanisms underlying inflammation in neurodegeneration. Cell, 2010. 140(6): p. 918–34.

57. Brambilla, R., et al., Inhibition of astroglial nuclear factor kappaB reduces inflammation and improves functional recovery after spinal cord injury. J Exp Med, 2005. 202(1): p. 145–56.

58. Zhang, C., et al., Cigarette smoke extract-induced p120-mediated NF-κB activation in human epithelial cells is dependent on the RhoA/ROCK pathway. Scientific Reports, 2016. 6(1): p. 23131.

59. Ganapathy, V., et al., Abstract 4470: E-cigarette aerosol exposure increases NF-kB and modulates inflammatory markers in oral epithelial cells. Cancer Research, 2023. 83(7_Supplement): p. 4470–4470.

60. Kauppinen, A., et al., Antagonistic crosstalk between NF-kappaB and SIRT1 in the regulation of inflammation and metabolic disorders. Cell Signal, 2013. 25(10): p. 1939–48.

61. Kauppinen, A., et al., Antagonistic crosstalk between NF-κB and SIRT1 in the regulation of inflammation and metabolic disorders. Cell Signal, 2013. 25(10): p. 1939–48.

62. Yeung, F., et al., Modulation of NF-kappaB-dependent transcription and cell survival by the SIRT1 deacetylase. Embo j, 2004. 23(12): p. 2369–80.

63. Yang, X.D., E. Tajkhorshid, and L.F. Chen, Functional interplay between acetylation and methylation of the RelA subunit of NF-kappaB. Mol Cell Biol, 2010. 30(9): p. 2170–80.

64. Durão, A.C.C.d.S., et al., In Utero Exposure to Environmental Tobacco Smoke Increases Neuroinflammation in Offspring. Frontiers in Toxicology, 2022. 3.

65. Nakahata, Y., et al., The NAD+-dependent deacetylase SIRT1 modulates CLOCK-mediated chromatin remodeling and circadian control. Cell, 2008. 134(2): p. 329–40.

66. Lagouge, M., et al., Resveratrol improves mitochondrial function and protects against metabolic disease by activating SIRT1 and PGC-1alpha. Cell, 2006. 127(6): p. 1109–22.

67. Rodgers, J.T., et al., Nutrient control of glucose homeostasis through a complex of PGC-1alpha and SIRT1. Nature, 2005. 434(7029): p. 113–8.

68. Picard, F., et al., Sirt1 promotes fat mobilization in white adipocytes by repressing PPAR-gamma. Nature, 2004. 429(6993): p. 771–6.

69. Bishop, N.A. and L. Guarente, Genetic links between diet and lifespan: shared mechanisms from yeast to humans. Nat Rev Genet, 2007. 8(11): p. 835–44.

70. Wu, W., et al., Nuclear factor-kappaB regulates the transcription of NADPH oxidase 1 in human alveolar epithelial cells. BMC Pulmonary Medicine, 2021. 21(1): p. 98.

71. Brunet, A., et al., Stress-dependent regulation of FOXO transcription factors by the SIRT1 deacetylase. Science, 2004. 303(5666): p. 2011–5.

72. Luo, J., et al., Negative control of p53 by Sir2alpha promotes cell survival under stress. Cell, 2001. 107(2): p. 137–48.

73. Yang, Y., et al., Regulation of SIRT1 and Its Roles in Inflammation. Front Immunol, 2022. 13: p. 831168.

74. Askin, L., et al., The relationship between coronary artery disease and SIRT1 protein. North Clin Istanb, 2020. 7(6): p. 631–635.

75. Mehramiz, M., et al., A Potential Role for Sirtuin-1 in Alzheimer’s Disease: Reviewing the Biological and Environmental Evidence. J Alzheimers Dis Rep, 2023. 7(1): p. 823–843.

76. Shinton, R. and G. Beevers, Meta-analysis of relation between cigarette smoking and stroke. BMJ, 1989. 298(6676): p. 789–94.

77. Wells, A.C. and S. Lotfipour, Prenatal nicotine exposure during pregnancy results in adverse neurodevelopmental alterations and neurobehavioral deficits. Advances in Drug and Alcohol Research, 2023. 3.

78. Hernández-Martínez, C., et al., Effects of Prenatal Nicotine Exposure on Infant Language Development: A Cohort Follow Up Study. Matern Child Health J, 2017. 21(4): p. 734–744.

79. Holbrook, B.D., The effects of nicotine on human fetal development. Birth Defects Res C Embryo Today, 2016. 108(2): p. 181–92.

80. Castro, E.M., S. Lotfipour, and F.M. Leslie, Nicotine on the developing brain. Pharmacological Research, 2023. 190: p. 106716.

81. Csabai, D., et al., Low intensity, long term exposure to tobacco smoke inhibits hippocampal neurogenesis in adult mice. Behav Brain Res, 2016. 302: p. 44–52.

82. Mills, A., et al., Influence of gestational window on offspring vascular health in rodents with in utero exposure to electronic cigarettes. J Physiol, 2024. 602(17): p. 4271–4289.

